# The fast-slow continuum is not the end-game of life history evolution, human or otherwise

**DOI:** 10.1101/2021.03.09.434595

**Authors:** Roberto Salguero-Gómez

## Abstract

Life history human evolution places a strong emphasis on the fast-slow continuum and pace-of-life continuum. Papers in this area frequently make a call for more integrative research to explore the relationships between the fast-slow continuum (population-level variation in traits) and pace-of-life continuum (ditto at the individual level). Here, using high-resolution demographic models from natural populations of 41 modern human populations, and hundreds of non-human animal populations and plant populations, I show that the fast-slow continuum, while of interest to explore variation in human strategies, should not be the sole focus of attention. Using a phylogenetically corrected principal component analysis, I show that other axes of life history variation show great promise. Specifically, understanding of the mechanisms of inter-specific and intra-specific differences in senescence rates in reproduction and survival, and how species respond to stochastic variation, I argue, should become the focus of my interdisciplinary research. The proposed new axis of demographic buffering-lability emerges as an important one to explain over 20% of variation in human and non-human populations. I show that these axes help predict demographic resilience, and also correlates with human nation per capita GDP and population growth rates.

---

Our understanding of nature is grounded in classification and comparison. By putting entities (e.g. individuals, populations, species) in boxes, by giving those boxes names, and then comparing and contrasting their attributes, we not only gain key insights into the wealth of diversity supported on Earth, but also into the evolutionary forces that have shaped it. Life history evolution is a branch of biology dedicated to understanding the diversity of life history traits (*i.e.* key moments of the life cycle of a species; including age at maturity, reproductive frequency, *etc.*) and life history strategies (*i.e.* combinations of life history traits that define the way of “making a living” of individuals, populations, or species; including long-lived semelparity, as in the chinook salmon [*Oncorhynchus tshawytscha*] or the century plant [*Agave americana*]). The main thesis of this commentary is that there is much to be gained in our understanding of human life histories by looking beyond the fast-slow continuum.

Life history theory dates from the very inception of ecology and evolution. Seminal works by evolutionary ecologists such as Pianka (1970), McArthur (1972), or Stearns (1992) continue to pave the way, several decades later, for our current vibrant research programme of life history theory. The Special Feature *“Current debates in human life history research”* is an excellent demonstration of this vibrancy. This special feature explores the drivers of life history strategies in humans, the scaling of strategies from individuals to population, and ways to best quantify trade-offs. The contributions provide key insights not only into anthropology and human psychology, but also into nearby disciplines such as ecology and evolution. I wish to highlight the wealth of cutting-edge research in non-human demography, not included in this special feature, which could drastically improve our understanding of (1) human life history evolution, (2) comparative demography, and (3) biodemography.

Much attention in life history theory has been paid to the so-called *fast-slow continuum.* All contributions to this special feature focus on this concept. In its original embodiment, this continuum is meant to organise the diversity of life history traits and strategies along a trade-off between large investment in survival (thus producing long-lived species, such as the killer whale [*Orcinus orca*] or the Eastern white pine [*Pinus strobus*]; see Figure 1A), on the one hand, and fast development and reproduction on the other (producing species with high generational turnover, such as the Queen Alexandra’s sulphur [*Colias Alexandra*] or the burnt orchid [*Neotinea ustulata*]). The applicability of this continuum to the pace-of-life continuum, which focuses on ranking individuals within a population, has been the focus of much research not only in humans (Willem & Nettle 2020), but also other species (Araya-Ajoy 2018).

**Figure 1.**
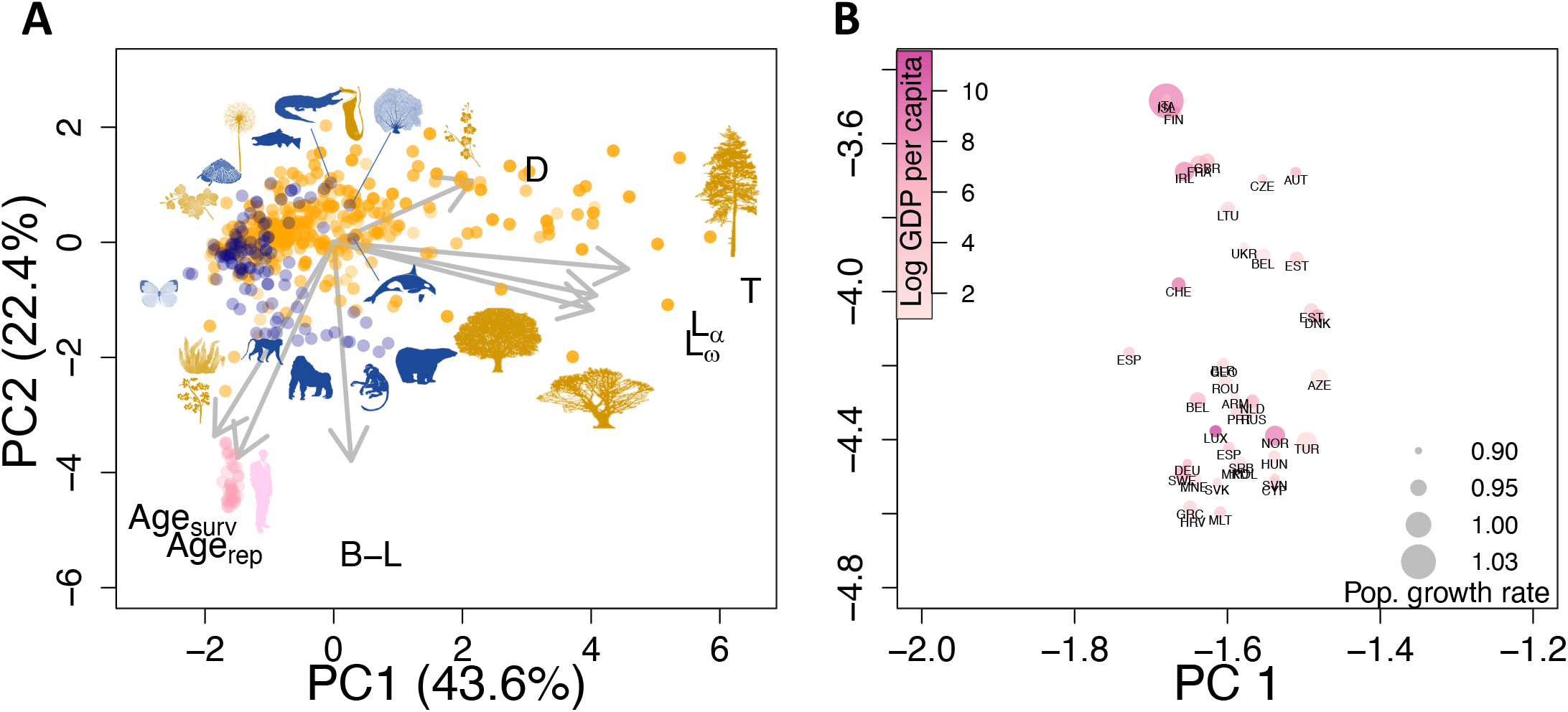
Human populations display a distinct set of life history traits from other animals and plants. Their life history trait location is not driven primarily by differences in the fast-slow continuum (T, *L_α_, L_ω_*, see below) but rather by their abnormally high rates of actuarial (Age_surv_) and reproductive senescence, as well as their extreme ability to buffer demographically (B-L). **A**. Phylogenetically controlled principal component analysis of seven key life history traits that define the dynamics of 41 modern human populations (pink), 111 non-human animal species (blue), and 784 plant populations (orange). The life history traits are: *T*: generation time; *L_α_:* age at maturity; *L_ω_:* mature lifespan; D: degree of parity; B-L: degree of demographic buffering-lability; Age_rep_: rate of reproductive senescence; and Age_surv_: rate of survival (actuarial) senescence. Silhouettes correspond to the following species (clock-wise order starting from the top right): Eastern white pine *(Pinus strobus),* smooth mesquite *(Prosopis laevigata),* sugar maple *(Acer saccharum),* killer whale *(Orcinus orca),* polar bear *(Ursus maritimus),* Northern muriqui *(Brachyteles hypoxanthus),* mountain gorilla *(Gorilla beringei beringei),* human *(Homo sapiens sapiens),* patas monkey *(Erythrocebus patas),* Himalayan blackberry *(Rubus praecox),* brown algae *(Cystoseira zosteroides),* Queen Alexandra’s sulphur *(Colias alexandra),* burnt orchid *(Neotinea ustulata),* deep sea limpet *(Lepetodrilus fucensis),* dandelion *(Taraxacum campylodes),* chinook salmon *(Oncorhynchus tshawytscha),* freshwater crocodile *(Crocodylus johnsoni),* pitcher plant *(Cirsium pitcheri),* red gorgonian *(aramuricea clavata),* and lady spider orchid *(Orchis purpurea).* **B**. A zoom-in to the life history trait space occupied by humans depicted in panel A, displaying the ISO3 country codes of the 41 examined nations, colour-coded by their per capita GDP, and with dot sizes proportional to their respective population growth rate trend in the last 30 years. Italy, Iceland and Finland are found at the top, while Malta, Greece and Croatia are at bottom, corresponding to populations with high senescence rates and high ability to buffer demographically. See Table S1 for further details.

The last decades have shed evidence on a much more complex picture of diversity of life history traits and strategies. Using multivariate analyses and large volumes of demographic data, Gaillard (1989), Salguero-Gómez et al. (2016), and Capdevila et al. (2020a), among others, demonstrated that the investment of energy into different moments of reproduction (e.g. age at first reproduction, reproductive window, degree of parity, annual intensity of reproduction) are decoupled from the fast-slow continuum in animals—humans included—and plants. Importantly, while the fast-slow continuum in these studies explains ~35% of the variation in life histories, the parity continuum explains ~30% in them. In a recent analysis contrasting the life history strategies of terrestrial *vs*. aquatic species, the parity axes actually gained more relevance than the fast-slow continuum (Capdevila et al. 2020a).

A main goal of life history and demography is to predict how individuals, populations, and species will respond to environmental stochasticity. This goal has recently become more pressing due to on-going climate change, increasing frequencies of disturbances (e.g. fires, pandemics, etc.). However, although links have been developed between life history continua and environmental stochasticity (Tuljapurkar et al. 2009), neither the fast-slow continuum nor the parity continuum explicitly address how life history traits respond to environmental stochasticity.

Over two decades ago, Pfister (1998) tackled this limitation in life history theory by introducing the demographic buffering hypothesis (DBH, hereafter). The DBH states that natural populations should regulate the temporal variation in their vital rates (e.g. survival, reproduction, development) to minimise the impact of environmental stochasticity. Based on *Tulja’s small noise approximation*, it is expected that the temporal variation of the vital rates that are more important for population growth rate *λ* (as quantified via sensitivity analysis) are constrained more than those with lower sensitivity. This is because the temporal autocorrelation of highly sensitive vital rates can bring down the stochastic population growth rate (λ_s_). Evidence of demographic buffering has been reported in some human populations, gorillas, and plants (see references in Hilde et al. 2020). However, numerous reports have recently emerged identifying an opposite demographic strategy that maximises λ_s_ in highly stochastic environments (references in Hilde et al. 2020). This strategy, the demographic lability hypothesis (DLH, hereafter), consists of investing in the vital rate that most matters to *λ* in years of plentiful resources, and not at all during bad environmental years. The booms experienced by the population during years of bonanza are then expected to outweigh the bursts that follow years of poor environmental quality for the demographic lability strategy to be adaptive. Though the expectation is that buffering populations are slow and labile populations fast (Hilde et al. 2020), evidence is starting to show that the fast-slow continuum axis is in fact orthogonal to whether populations buffer or act labilly to the environment. As a consequence, here I propose that a third axes of life history variation exists: the buffering-lability axis.

Three life history continua to rule them all? What utility emerges from expanding the toolbox of demographers (humans and biodemographers alike) along three axes of variation? In a multivariate analysis of the life history traits of 111 animals (including humans) and 784 plant species (Figure 1A), I show that three axes are necessary to explain 80% of variance. PC1 explains 43.6% of the variance and corresponds to the fast-slow continuum. PC2 explains 22.4% and is predominantly explained by senescence and demographic buffering-lability. PC3 (not shown) explains 13.3% of the variance and corresponds to the degree of parity. This finding contrasts with the original works on life history, which suggested that the fast-slow continuum alone should explain this large degree of variation (Stearns 1992). Moreover, the usage of the parity continuum allows us to closely examine non-decaying functions. The fast-slow continuum is mostly driven by an ever-declining function: survival. However, reproduction can take a myriad of shapes, with some populations having a sharp increase followed by a sharp decline (e.g. salmons, agaves), and others having frequent bouts of reproduction with breaks in between (e.g. masting as in oaks [*Quercus spp*.]). With the diversity of shapes that reproduction can take, important moments emerge that are not accounted for by the fast-slow continuum, including the frequency of reproduction (*D*) or reproductive senescence *(Age_rep_*). Furthermore, the buffering-lability axis adds a new, valuable perspective not explicitly considered by the fast-slow continuum: as the investment of energy into survival, development, reproduction, and recruitment can vary through time, so can the life history traits and strategies that emanate from said decisions. When compared to their closely related animal species, and to the plant kingdom, humans do not actually vary much along the fast slow continuum, but rather along a continuum of actuarial and reproductive senescence, and along the extreme of demographic buffering (Figure 1A). In this case, the 41 examined human populations have a high sensitivity to adult survival, and their survival rates have not changed much in the last decades - so humans are extreme demographic “bufferers”.

When examined closely, important predictions can be drawn from this senescence/buffering continuum in human populations. The ranking of human populations along this continuum of senescence and demographic buffering-lability predicts key properties. Using the scores of the 41 human populations as indicators of their placing along PC1 (fast-slow continuum) and PC2 (senescence/buffering-using lability continuum), we can now ask whether these help predict key demographic and socio-economic properties. Both axes are significantly positively correlated with the ability of those populations to quickly recover from disturbances, as quantified by the damping ratio (Capdevila et al. 2020b) of the respective matrix population models (PC1: *P*<0.031; PC2: *P*<0.032). While PC1 and PC2 do not predict the rate of growth of the population, PC3 is positively correlated with the growth rate (*P*<0.002), such that human populations with higher reproductive frequencies grow faster. Similarly, both PC1 and PC2 are significantly negatively correlated with the per capita GDP of the country (PC1: *P*<0.030; PC2: *P*<0.028, Figure 1B). The examination of how different behaviours accelerate the rates of actuarial and reproductive senescence is not new in (human) biology (Jones et al. 2004), but it was an aspect that papers in this special feature did not cover. Investigations of how individual behaviours may range within the same human population from more demographically buffering (or *homeostatic,* see Hilde et al. 2020) to more demographically labile (or variable), I argue, holds great promise to link human evolutionary and behavioural studies.

Virtually all papers in this special feature, including the editorial (Frankenhuis & Nettle 2020), make a call for more collaborative research between life history research in psychology and life history research in biology. Here, using big data (*i.e.,* 941 species, including multiple populations of modern humans, other animals, and plants), I have shown that important axes of life history trait variation remain overlooked in human life history research. While exploring the drivers of said variation is crucial, it is also key to note that we may learn much more about humans by examining species that are much closer to us in the life history trait space (e.g. Himalayan blackberry *[Rubus praecox])* than phylogenetically (e.g. all primates are rather far from humans in Figure 1A). I encourage researchers interested in these kinds of questions to examine whether/how these continua shape the behaviour and evolution of humans, and to take inspiration from the comparative approach employed here to do so.

## Acknowledgements

I thank T. Ezard for making the human demographic models open-access, as well as the hundreds of population ecologists who have deposited their animal and plant demographic models in www.compadre-db.org. Due to reference limitations, a full list of the citations used in Figure 1 can be found in Table S1. This work was supported by a NERC Independent Research Fellowship (NE/M018458/1). I thank K. Davis for comments to improve the readability of this piece.

