## Supplementary Materials Table 1 and 2 for "The fast-slow continuum is not the end-game of life history evolution, human or otherwise"

Roberto Salguero-Gómez

**Table S1**. Summary of data sources from which life history traits were derived using age-from-stage decompositions on matrix population models (Caswell 2001) stored in the COMADRE Animal Matrix Database v. 4.21.1.0 (Salguero-Gómez et al. 2016a) and COMPADRE Plant Matrix Database v. 4.21.1.0 (Salguero-Gómez et al. 2015). Briefly, the life history traits that I derived from this matrix population models are: *T*: Generation time (years); *L_α_*: age at maturity (years); *L_ω_*; mature lifespan (years); *D*: Demetrius entropy (Salguero-Gómez et al. 2016b) or degree of parity (dimensionless). *D* quantifies the degree of parity, with strictly semelparous species (only one reproductive bout) having a value of 1, and increasing values with the degree of iteroparity; *B-L*: degree of demographic buffering-lability (dimensionless). The B-L values are the correlation coefficient of the variance in vital rates and the sensitivity of population growth rate to those vital rates as per Pfister (1998); *Age_Rep_*: rate of reproductive senescence (dimensionless), as detailed in Baudisch and Stott (2019); *Age_Surv_*: rate of actuarial senescence (dimensionless), following the same logic as *Age_Rep_*. These life history traits were obtained using the Rage R library (<https://github.com/jonesor/Rage>).

| **Species** | **Kingdom** | **Authors** | **Journal** | **Year** | **DOI_ISBN** |
| --- | --- | --- | --- | --- | --- |
| *Alouatta seniculus* | Animalia | Wiederholt; Fernandez-Duque; Diefenbach; Rudran | Ecol Model | 2010 | 10.1016/j.ecolmodel.2010.06.026 |
| *Brachyteles hypoxanthus* | Animalia | Morris; Pfister; Tuljapurkar; Haridas; Boggs; Boyce; Bruna; Church; Coulson; Doak; Forsyth; Gaillard; Horvitz; Kalisz; Kendall; Knight; Lee; Menges | Ecology | 2008 | 10.1890/07-0774.1 |
| *Cebus capucinus* | Animalia | Morris; Pfister; Tuljapurkar; Haridas; Boggs; Boyce; Bruna; Church; Coulson; Doak; Forsyth; Gaillard; Horvitz; Kalisz; Kendall; Knight; Lee; Menges | Ecology | 2008 | 10.1890/07-0774.1 |
| *Cervus canadensis* | Animalia | Clark | NA | 2014 | NA |
| *Chlorocebus aethiops* | Animalia | Isbell; Young; Jaffe; Carlson; Chancellor | Int J Primatol | 2009 | 10.1007/s10764-009-9332-7 |
| *Dasypus novemcinctus* | Animalia | Oli; Loughry; Caswell; Perez-Heydrich; McDonough; Truman | Ecol Model | 2017 | 10.1016/j.ecolmodel.2017.02.001 |
| *Erythrocebus patas* | Animalia | Isbell; Young; Jaffe; Carlson; Chancellor | Int J Primatol | 2009 | 10.1007/s10764-009-9332-7 |
| *Giraffa camelopardalis* | Animalia | Strauss; Kilewo; Rentsch; Packer | Popul Ecol | 2015 | 10.1007/s10144-015-0499-9 |
| *Gorilla beringei beringei* | Animalia | Morris; Pfister; Tuljapurkar; Haridas; Boggs; Boyce; Bruna; Church; Coulson; Doak; Forsyth; Gaillard; Horvitz; Kalisz; Kendall; Knight; Lee; Menges | Ecology | 2008 | 10.1890/07-0774.1 |
| *Ovis canadensis* | Animalia | Coulson; Gaillard; Festa-Bianchet | J Anim Ecol | 2005 | 10.1111/j.1365-2656.2005.00975.x |
| *Puma concolor* | Animalia | Lambert; Wielgus; Robinson; Katnik; Cruickshank; Clarke; Almack | J Wildlife Manage | 2006 | 10.2193/0022-541X(2006)70[246:CPDAVI]2.0.CO;2 |
| *Pygoscelis adeliae* | Animalia | Hinke; Trivelpiece; Trivelpiece | Ecosphere | 2017 | 10.1002/ecs2.1666 |
| *Zalophus californianus* | Animalia | Wielgus; Gonzalez-Suarez; Aurioles-Gamboa; Gerber | Ecol Appl | 2008 | 10.1890/07-0892.1 |
| *Rana temporaria* | Animalia | Campbell; Garner; Tessa; Scheele; Griffiths; Wilfert; Harrison | PeerJ | 2018 | NA |
| *Astroblepus ubidiai* | Animalia | V√©lez-Espino | Ecol Freshw Fish | 2005 | 10.1111/j.1600-0633.2005.00084.x |
| *Clinocottus analis* | Animalia | Davis; Levin | Mar Ecol Prog Ser | 2002 | 10.3354/meps234229 |
| *Clinostomus funduloides* | Animalia | Peoples | Master Thesis | 2010 | NA |
| *Cottus bairdi* | Animalia | Peoples | Master Thesis | 2010 | NA |
| *Cyprinodon diabolis* | Animalia | Beissinger | PeerJ | 2014 | 10.7717/peerj.549 |
| *Oncorhynchus tshawytscha* | Animalia | Wilson | Conserv Biol | 2003 | 10.1046/j.1523-1739.2003.01535.x |
| *Salvelinus confluentus* | Animalia | Bowerman | NA | 2013 | NA |
| *Zingel asper* | Animalia | Labonne; Gaudin | Can J Fish Aquat Sci | 2006 | 10.1139/f05-245 |
| *Zoarces viviparus* | Animalia | Bergek; Ma; Vetemaa; Franz√©n; Appelberg | Ecotox Environ Safe | 2012 | 10.1016/j.ecoenv.2012.01.019 |
| *Geocrinia alba* | Animalia | Conroy; Brook | Popul Ecol | 2003 | 10.1007/s10144-003-0145-9 |
| *Geocrinia vitellina* | Animalia | Conroy; Brook | Popul Ecol | 2003 | 10.1007/s10144-003-0145-9 |
| *Paramuricea clavata* | Animalia | Linares; Doak | Mar Ecol Prog Ser | 2010 | 10.3354/meps08437 |
| *Paramuricea clavata* | Animalia | Linares; Doak; Coma; Diaz; Zabala | Ecology | 2007 | 10.1890/05-1931 |
| *Anas laysanensis* | Animalia | Reynolds; Weiser; Jamieson; Hatfield | J Wildlife Manage | 2013 | 10.1002/jwmg.582 |
| *Chen caerulescens* | Animalia | Cooch; Rockwell; Brault | Ecol Monogr | 2001 | 10.1890/0012-9615(2001)071[0377:RAODRT]2.0.CO;2 |
| *Anthropoides paradiseus* | Animalia | Altwegg; Anderson | Funct Ecol | 2009 | 10.1111/j.1365-2435.2009.01563.x |
| *Bonasa umbellus* | Animalia | Tirpak; Giuliano; Miller; Allen; Bittner; Buehler; Edwards; Harper; Igo; Norman; Seamster; Stauffer | Biol Conserv | 2006 | 10.1016/j.biocon.2006.06.014 |
| *Buteo solitarius* | Animalia | Klavitter; Marzluff; Vekasy | J Wildlife Manage | 2003 | 10.2307/3803072 |
| *Campylorhynchus brunneicapillus sandiegensis* | Animalia | Conlisk; Motheral; Chung; Wisinski; Endress | Biol Conserv | 2014 | 10.1016/j.biocon.2014.04.010 |
| *Centrocercus minimus* | Animalia | Davis; Hooten; Phillips; Doherty | Ecol Evol | 2014 | 10.1002/ece3.1290 |
| *Ciconia ciconia* | Animalia | Schaub; Pradel; Lebreton | Biol Conserv | 2004 | 10.1016/j.biocon.2003.11.002 |
| *Falco naumanni* | Animalia | Hiraldo; Negro; Donazar; Gaona | J Appl Ecol | 1996 | 10.2307/2404688 |
| *Falco peregrinus* | Animalia | Altwegg; Jenkins; Abadi | Ibis | 2013 | 10.1111/ibi.12125 |
| *Forpus passerinus* | Animalia | Sandercock; Beissinger | J Appl Stat | 2002 | 10.1080/02664760120108818 |
| *Himantopus novaezelandiae* | Animalia | Cruz; Pech; Seddon; Cleland; Nelson; Sanders; Maloney | Biol Conserv | 2013 | 10.1016/j.biocon.2013.09.006 |
| *Lagopus leucura* | Animalia | Wilson; Martin | BMC Ecol | 2012 | 10.1186/1472-6785-12-9 |
| *Lagopus muta* | Animalia | Wilson; Martin | BMC Ecol | 2012 | 10.1186/1472-6785-12-9 |
| *Pernis apivorus* | Animalia | Bijlsma; Vermeulen; Hemerik; Klok | Ardea | 2012 | 10.5253/078.100.0208 |
| *Phalacrocorax auritus* | Animalia | Chastant; King; Weseloh; Moore | J Wildlife Manage | 2014 | 10.1002/jwmg.628 |
| *Sterna hirundo* | Animalia | Szostek; Schaub; Becker | J Anim Ecol | 2014 | 10.1111/1365-2656.12206 |
| *Strix occidentalis* | Animalia | LaHaye; Zimmerman; Guti√©rrez | Auk | 2004 | 10.1642/0004-8038(2004)121[1056:TVITVR]2.0.CO;2 |
| *Turdus torquatus* | Animalia | Sim; Rebecca; Ludwig; Grant; Reid | J Anim Ecol | 2011 | 10.1111/j.1365-2656.2010.01750.x |
| *Nuttallia obscurata* | Animalia | Dudas; Dower; Anholt | Ecology | 2007 | 10.1890/06-1216.1 |
| *Amphimedon compressa* | Animalia | Mercado-Molina; Sabat; Yoshioka | J Exp Mar Biol Ecol | 2011 | 10.1016/j.jembe.2011.07.018 |
| *Spongia graminea* | Animalia | Cropper; Di Resta | Ecol Model | 1999 | 10.1016/S0304-3800(99)00039-3 |
| *Xestospongia muta* | Animalia | McMurray; Henkel; Pawlik | Ecology | 2010 | 10.3354/meps339093 |
| *Lepetodrilus fucensis* | Animalia | Kelly; Metaxas | Mar Ecol Prog Ser | 2010 | 10.3354/meps08442 |
| *Umbonium costatum* | Animalia | Noda; Nakao | J Anim Ecol | 1996 | 10.2307/5722 |
| *Homo sapiens* | Animalia | Keyfitz; Flieger | NA | 1990 | 0-226-43237-8 |
| *Homo sapiens* | Animalia | Keyfitz; Flieger | NA | 1990 | 0-226-43237-8 |
| *Acyrthosiphon pisum* | Animalia | Gross; Craig; Hutchinson | Ecology | 2002 | 10.1890/0012-9658(2002)083[3285:BEOADM]2.0.CO;2 |
| *Cicindela ohlone* | Animalia | Cornelisse; Bennett; Letourneau | PLOS ONE | 2013 | 10.1371/journal.pone.0071005 |
| *Colias alexandra* | Animalia | Hayes | Oecologia | 1981 | 10.1007/BF00349187 |
| *Scolytus ventralis* | Animalia | Berryman | Can Entomol | 1973 | 10.4039/Ent1051465-11 |
| *Scolytus ventralis* | Animalia | Berryman | Can Entomol | 1973 | 10.4039/Ent1051465-11 |
| *Callinectes sapidus* | Animalia | Miller | Estuaries | 2001 | 10.2307/1353238 |
| *Antechinus agilis* | Animalia | Lindenmayer; Lacy | Biol Conserv | 2002 | 10.1016/S0006-3207(01)00134-3 |
| *Brachyteles hypoxanthus* | Animalia | Morris; Altmann; Brockman; Cords; Fedigan; Pusey; Stoinski; Bronikowski; Alberts; Strier | Am Nat | 2011 | 10.1086/657443 |
| *Callospermophilus lateralis* | Animalia | Hostetler; Kneip; Van Vuren; Oli | PLOS ONE | 2012 | 10.1371/jourNAl.pone.0034379 |
| *Cebus capucinus* | Animalia | Morris; Altmann; Brockman; Cords; Fedigan; Pusey; Stoinski; Bronikowski; Alberts; Strier | Am Nat | 2011 | 10.1086/657443 |
| *Cercopithecus mitis* | Animalia | Morris; Altmann; Brockman; Cords; Fedigan; Pusey; Stoinski; Bronikowski; Alberts; Strier | Am Nat | 2011 | 10.1086/657443 |
| *Clethrionomys rufocanus* | Animalia | Yoccoz; Nakata; Stenseth; Saitoh | Res Popul Ecol | 1998 | 10.1007/BF02765226 |
| *Eumetopias jubatus* | Animalia | Holmes; York | Conserv Biol | 2003 | 10.1111/j.1523-1739.2003.00191.x |
| *Felis catus* | Animalia | Budke; Slater | J Appl Anim Welf Sci | 2009 | 10.1080/10888700903163419 |
| *Gorilla beringei beringei* | Animalia | Morris; Altmann; Brockman; Cords; Fedigan; Pusey; Stoinski; Bronikowski; Alberts; Strier | Am Nat | 2011 | 10.1086/657443 |
| *Hippocamelus bisulcus* | Animalia | Corti; Wittmer; Festa-Bianchet | J Mammal | 2010 | 10.1644/09-MAMM-A-047.1 |
| *Macaca mulatta* | Animalia | Hern√°ndez-Pacheco; Rawlins; Kessler; Williams; Ruiz-Maldonado; Gonz√°lez-Martinez; Ruiz-Lambides; Sabat | Am J Primatol | 2013 | 10.1002/ajp.22177 |
| *Macaca mulatta* | Animalia | Kessler; Pacheco; Rawlings; Ruiz-Lambrides; Delgado; Sabat | Am J Primatol | 2014 | 10.1002/ajp.22323 |
| *Macropus eugenii* | Animalia | Chambers; Bencini | Wildlife Res | 2010 | 10.1071/WR10080 |
| *Marmota flaviventris* | Animalia | Ozgul; Oli; Armitage; Blumstein; Van Vuren | Am Nat | 2009 | 10.1086/597225 |
| *Marmota flaviventris* | Animalia | Ozgul; Oli; Armitage; Blumstein; Van Vuren | Am Nat | 2009 | 10.1086/597225 |
| *Microtus oeconomus* | Animalia | Johannesen; Aars; Andreassen; Ims | Popul Ecol | 2003 | 10.1007/s10144-003-0139-7 |
| *Mustela erminea* | Animalia | Wittmer; Powell; King | J Anim Ecol | 2007 | 10.1111/j.1365-2656.2007.01274.x |
| *Odocoileus virginianus* | Animalia | Chitwood; Lashley; Kilgo; Moorman; Deperno | J Wildlife Manage | 2015 | 10.1002/jwmg.835 |
| *Orcinus orca* | Animalia | V√©lez-Espino; Ford; Ara√∫jo; Ellis; Parken; Balcomb | Can Tech Report Fish & Aq Sci | 2014 | 978-1-100-23563-9 |
| *Ovis aries* | Animalia | Clutton-Brock; Price; Albon; Jewell | J Anim Ecol | 1992 | 10.2307/5330 |
| *Ovis canadensis* | Animalia | Rubin; Boyce; Caswell-Chen | J Wildlife Manage | 2002 | 10.2307/3803144 |
| *Ovis canadensis* | Animalia | Johnson; Mills; Wehausen; Stephenson | Ecology | 2010 | 10.1111/j.1365-2664.2010.01846.x |
| *Pan troglodytes schweinfurthii* | Animalia | Morris; Altmann; Brockman; Cords; Fedigan; Pusey; Stoinski; Bronikowski; Alberts; Strier | Am Nat | 2011 | 10.1086/657443 |
| *Papio cynocephalus* | Animalia | Morris; Altmann; Brockman; Cords; Fedigan; Pusey; Stoinski; Bronikowski; Alberts; Strier | Am Nat | 2011 | 10.1086/657443 |
| *Phocarctos hookeri* | Animalia | Lalas; Bradshaw | Biol Conserv | 2003 | 10.1016/S0006-3207(02)00421-4 |
| *Propithecus verreauxi* | Animalia | Morris; Altmann; Brockman; Cords; Fedigan; Pusey; Stoinski; Bronikowski; Alberts; Strier | Am Nat | 2011 | 10.1086/657443 |
| *Puma concolor* | Animalia | Lambert; Wielgus; Robinson; Katnik; Cruickshank; Clarke; Almack | J Wildlife Manage | 2006 | 10.2193/0022-541X(2006)70[246:CPDAVI]2.0.CO;2 |
| *Rattus fuscipes* | Animalia | Lindenmayer; Lacy | Biol Conserv | 2002 | 10.1016/S0006-3207(01)00134-3 |
| *Saguinus fuscicollis* | Animalia | Watsa | NA | 2013 | 10.7936/K7DB7ZTD |
| *Sigmodon hispidus* | Animalia | Sauer; Slade | J Mammal | 1985 | 10.2307/1381244 |
| *Sigmodon hispidus* | Animalia | Sauer; Slade | J Mammal | 1985 | 10.2307/1381244 |
| *Urocitellus armatus* | Animalia | Oli; Slade; Dobson | Ecology | 2001 | 10.1890/0012-9658(2001)082[1921:EODROU]2.0.CO;2 |
| *Urocitellus armatus* | Animalia | Oli; Slade; Dobson | Ecology | 2001 | 10.1890/0012-9658(2001)082[1921:EODROU]2.0.CO;2 |
| *Ursus arctos* | Animalia | Wielgus | Biol Conserv | 2002 | 10.1016/S0006-3207(01)00265-8 |
| *Ursus maritimus* | Animalia | Hunter; Caswell; Runge; Regehr; Amstrup; Stirling | Ecology | 2010 | 10.1890/09-1641 |
| *Arctodiaptomus salinus* | Animalia | Jim√©nez-Melero; Ramirez; Guerrero | Freshwater Biol | 2013 | 10.3354/meps10377 |
| *Apalone mutica* | Animalia | Zimmer-Shaffer; Briggler; Millspaugh | Chelonian Conserv Bi | 2014 | 10.2744/CCB-1109.1 |
| *Apalone spinifera* | Animalia | Zimmer-Shaffer; Briggler; Millspaugh | Chelonian Conserv Bi | 2014 | 10.2744/CCB-1109.1 |
| *Chelydra serpentina* | Animalia | Zimmer-Shaffer; Briggler; Millspaugh | Chelonian Conserv Bi | 2014 | 10.2744/CCB-1109.1 |
| *Chrysemys picta* | Animalia | Mitchell | Herpetol Monogr | 1988 | 10.2307/1467026 |
| *Crocodylus acutus* | Animalia | Richards | Biological Sciences | 2003 | NA |
| *Crocodylus johnsoni* | Animalia | Tucker | NA | 2001 | 978-0-949324-89-4 |
| *Kinosternon integrum* | Animalia | Macip-R√≠os; Brauer-Robleda; Z√∫√±iga-Vega; Casas-Andreu | Herpetol J | 2011 | NA |
| *Kinosternon subrubrum* | Animalia | Frazer; Gibbons; Greene | Ecology | 1991 | 10.2307/1941572 |
| *Sceloporus arenicolus* | Animalia | Ryberg; Hill; Painter; Fitzgerald | Conserv Biol | 2014 | 10.1111/cobi.12429 |
| *Sceloporus grammicus* | Animalia | P√©rez-Mendoza | Herpetologica | 2013 | 10.1655/HERPETOLOGICA-D-12-00038R2 |
| *Sceloporus grammicus* | Animalia | M√©ndez‚Äìde la Cruz; Z√∫√±iga-Vega; Cuellar | Can J Zool | 2008 | 10.1139/Z08-124 |
| *Xenosaurus grandis* | Animalia | Z√∫√±iga-Vega; Valverde; Rojas-Gonzalez; Lemos-Espinal | Copeia | 2007 | 10.1643/0045-8511(2007)7[324:AOTPDO]2.0.CO;2 |
| *Xenosaurus platyceps* | Animalia | Rojas-Gonzalez; Jones; Z√∫√±iga-Vega; Lemos-Espinal | Amphibia-Reptilia | 2008 | 10.1163/156853808784124992 |
| *Xenosaurus sp.* | Animalia | Zamora-Abrego; Chang; Z√∫√±iga-Vega; Nieto-Montes de Oca; Johnson | Herpetologica | 2010 | 10.1655/09-005.1 |
| *Homo sapiens sapiens* | Animalia | Nicol-Harper; Dooley; Packman; Mueller; Bijak; Hodgson; Townley; Ezard | Popul Ecol | 2018 | 10.1007/s10144-018-0620-y |
| *Homo sapiens sapiens* | Animalia | Nicol-Harper; Dooley; Packman; Mueller; Bijak; Hodgson; Townley; Ezard | Popul Ecol | 2018 | 10.1007/s10144-018-0620-y |
| *Homo sapiens sapiens* | Animalia | Nicol-Harper; Dooley; Packman; Mueller; Bijak; Hodgson; Townley; Ezard | Popul Ecol | 2018 | 10.1007/s10144-018-0620-y |
| *Homo sapiens sapiens* | Animalia | Nicol-Harper; Dooley; Packman; Mueller; Bijak; Hodgson; Townley; Ezard | Popul Ecol | 2018 | 10.1007/s10144-018-0620-y |
| *Homo sapiens sapiens* | Animalia | Nicol-Harper; Dooley; Packman; Mueller; Bijak; Hodgson; Townley; Ezard | Popul Ecol | 2018 | 10.1007/s10144-018-0620-y |
| *Homo sapiens sapiens* | Animalia | Nicol-Harper; Dooley; Packman; Mueller; Bijak; Hodgson; Townley; Ezard | Popul Ecol | 2018 | 10.1007/s10144-018-0620-y |
| *Homo sapiens sapiens* | Animalia | Nicol-Harper; Dooley; Packman; Mueller; Bijak; Hodgson; Townley; Ezard | Popul Ecol | 2018 | 10.1007/s10144-018-0620-y |
| *Homo sapiens sapiens* | Animalia | Nicol-Harper; Dooley; Packman; Mueller; Bijak; Hodgson; Townley; Ezard | Popul Ecol | 2018 | 10.1007/s10144-018-0620-y |
| *Homo sapiens sapiens* | Animalia | Nicol-Harper; Dooley; Packman; Mueller; Bijak; Hodgson; Townley; Ezard | Popul Ecol | 2018 | 10.1007/s10144-018-0620-y |
| *Homo sapiens sapiens* | Animalia | Nicol-Harper; Dooley; Packman; Mueller; Bijak; Hodgson; Townley; Ezard | Popul Ecol | 2018 | 10.1007/s10144-018-0620-y |
| *Homo sapiens sapiens* | Animalia | Nicol-Harper; Dooley; Packman; Mueller; Bijak; Hodgson; Townley; Ezard | Popul Ecol | 2018 | 10.1007/s10144-018-0620-y |
| *Homo sapiens sapiens* | Animalia | Nicol-Harper; Dooley; Packman; Mueller; Bijak; Hodgson; Townley; Ezard | Popul Ecol | 2018 | 10.1007/s10144-018-0620-y |
| *Homo sapiens sapiens* | Animalia | Nicol-Harper; Dooley; Packman; Mueller; Bijak; Hodgson; Townley; Ezard | Popul Ecol | 2018 | 10.1007/s10144-018-0620-y |
| *Homo sapiens sapiens* | Animalia | Nicol-Harper; Dooley; Packman; Mueller; Bijak; Hodgson; Townley; Ezard | Popul Ecol | 2018 | 10.1007/s10144-018-0620-y |
| *Homo sapiens sapiens* | Animalia | Nicol-Harper; Dooley; Packman; Mueller; Bijak; Hodgson; Townley; Ezard | Popul Ecol | 2018 | 10.1007/s10144-018-0620-y |
| *Homo sapiens sapiens* | Animalia | Nicol-Harper; Dooley; Packman; Mueller; Bijak; Hodgson; Townley; Ezard | Popul Ecol | 2018 | 10.1007/s10144-018-0620-y |
| *Homo sapiens sapiens* | Animalia | Nicol-Harper; Dooley; Packman; Mueller; Bijak; Hodgson; Townley; Ezard | Popul Ecol | 2018 | 10.1007/s10144-018-0620-y |
| *Homo sapiens sapiens* | Animalia | Nicol-Harper; Dooley; Packman; Mueller; Bijak; Hodgson; Townley; Ezard | Popul Ecol | 2018 | 10.1007/s10144-018-0620-y |
| *Homo sapiens sapiens* | Animalia | Nicol-Harper; Dooley; Packman; Mueller; Bijak; Hodgson; Townley; Ezard | Popul Ecol | 2018 | 10.1007/s10144-018-0620-y |
| *Homo sapiens sapiens* | Animalia | Nicol-Harper; Dooley; Packman; Mueller; Bijak; Hodgson; Townley; Ezard | Popul Ecol | 2018 | 10.1007/s10144-018-0620-y |
| *Homo sapiens sapiens* | Animalia | Nicol-Harper; Dooley; Packman; Mueller; Bijak; Hodgson; Townley; Ezard | Popul Ecol | 2018 | 10.1007/s10144-018-0620-y |
| *Homo sapiens sapiens* | Animalia | Nicol-Harper; Dooley; Packman; Mueller; Bijak; Hodgson; Townley; Ezard | Popul Ecol | 2018 | 10.1007/s10144-018-0620-y |
| *Homo sapiens sapiens* | Animalia | Nicol-Harper; Dooley; Packman; Mueller; Bijak; Hodgson; Townley; Ezard | Popul Ecol | 2018 | 10.1007/s10144-018-0620-y |
| *Homo sapiens sapiens* | Animalia | Nicol-Harper; Dooley; Packman; Mueller; Bijak; Hodgson; Townley; Ezard | Popul Ecol | 2018 | 10.1007/s10144-018-0620-y |
| *Homo sapiens sapiens* | Animalia | Nicol-Harper; Dooley; Packman; Mueller; Bijak; Hodgson; Townley; Ezard | Popul Ecol | 2018 | 10.1007/s10144-018-0620-y |
| *Homo sapiens sapiens* | Animalia | Nicol-Harper; Dooley; Packman; Mueller; Bijak; Hodgson; Townley; Ezard | Popul Ecol | 2018 | 10.1007/s10144-018-0620-y |
| *Homo sapiens sapiens* | Animalia | Nicol-Harper; Dooley; Packman; Mueller; Bijak; Hodgson; Townley; Ezard | Popul Ecol | 2018 | 10.1007/s10144-018-0620-y |
| *Homo sapiens sapiens* | Animalia | Nicol-Harper; Dooley; Packman; Mueller; Bijak; Hodgson; Townley; Ezard | Popul Ecol | 2018 | 10.1007/s10144-018-0620-y |
| *Homo sapiens sapiens* | Animalia | Nicol-Harper; Dooley; Packman; Mueller; Bijak; Hodgson; Townley; Ezard | Popul Ecol | 2018 | 10.1007/s10144-018-0620-y |
| *Homo sapiens sapiens* | Animalia | Nicol-Harper; Dooley; Packman; Mueller; Bijak; Hodgson; Townley; Ezard | Popul Ecol | 2018 | 10.1007/s10144-018-0620-y |
| *Homo sapiens sapiens* | Animalia | Nicol-Harper; Dooley; Packman; Mueller; Bijak; Hodgson; Townley; Ezard | Popul Ecol | 2018 | 10.1007/s10144-018-0620-y |
| *Homo sapiens sapiens* | Animalia | Nicol-Harper; Dooley; Packman; Mueller; Bijak; Hodgson; Townley; Ezard | Popul Ecol | 2018 | 10.1007/s10144-018-0620-y |
| *Homo sapiens sapiens* | Animalia | Nicol-Harper; Dooley; Packman; Mueller; Bijak; Hodgson; Townley; Ezard | Popul Ecol | 2018 | 10.1007/s10144-018-0620-y |
| *Homo sapiens sapiens* | Animalia | Nicol-Harper; Dooley; Packman; Mueller; Bijak; Hodgson; Townley; Ezard | Popul Ecol | 2018 | 10.1007/s10144-018-0620-y |
| *Homo sapiens sapiens* | Animalia | Nicol-Harper; Dooley; Packman; Mueller; Bijak; Hodgson; Townley; Ezard | Popul Ecol | 2018 | 10.1007/s10144-018-0620-y |
| *Homo sapiens sapiens* | Animalia | Nicol-Harper; Dooley; Packman; Mueller; Bijak; Hodgson; Townley; Ezard | Popul Ecol | 2018 | 10.1007/s10144-018-0620-y |
| *Homo sapiens sapiens* | Animalia | Nicol-Harper; Dooley; Packman; Mueller; Bijak; Hodgson; Townley; Ezard | Popul Ecol | 2018 | 10.1007/s10144-018-0620-y |
| *Homo sapiens sapiens* | Animalia | Nicol-Harper; Dooley; Packman; Mueller; Bijak; Hodgson; Townley; Ezard | Popul Ecol | 2018 | 10.1007/s10144-018-0620-y |
| *Homo sapiens sapiens* | Animalia | Nicol-Harper; Dooley; Packman; Mueller; Bijak; Hodgson; Townley; Ezard | Popul Ecol | 2018 | 10.1007/s10144-018-0620-y |
| *Homo sapiens sapiens* | Animalia | Nicol-Harper; Dooley; Packman; Mueller; Bijak; Hodgson; Townley; Ezard | Popul Ecol | 2018 | 10.1007/s10144-018-0620-y |
| *Ascophyllum nodosum* | Chromalveolata | Aberg | Mar Ecol Prog Ser | 1990 | 10.3354/meps063281 |
| *Astragalus cottonii* | Plantae | Kaye | NA | 1990 | NA |
| *Astragalus bibullatus* | Plantae | Albrecht; Knight; Bernardo | Biol Conserv | 2016 | 10.1016/j.biocon.2016.09.030 |
| *Chamaedorea radicalis* | Plantae | Ash; Gorchov; Endress | Southwest Nat | 2013 | 10.1894/0038-4909-58.1.70 |
| *Cirsium pitcheri* | Plantae | Halsey; Bell; McEachern; Pavlovic | Ecosphere | 2016 | 10.1002/ecs2.1536 |
| *Conradina glabra* | Plantae | Bladow; Bohner; Winn | Bio One | 2017 | 10.3375/043.037.0305 |
| *Cyrtandra dentata* | Plantae | Bialic-Murphy; Gaoue; Kawelo | J Appl Ecol | 2017 | 10.1111/1365-2664.12868 |
| *Cystoseira zosteroides* | Plantae | Capdevila; Hereu; Riera; Linares | Ecology | 2016 | 10.1111/1365-2745.12625 |
| *Daphne rodriguezii* | Plantae | Rodriguez-Perez; Traveset | Oikos | 2012 | 10.1111/j.1600-0706.2011.19946.x |
| *Dracocephalum austriacum* | Plantae | Dost√°lek; M√ºnzbergov√° | Folia Geobot | 2013 | NA |
| *Dracocephalum austriacum* | Plantae | Dost√°lek; M√ºnzbergov√° | Folia Geobot | 2013 | NA |
| *Echinacea angustifolia* | Plantae | Dykstra | NA | 2013 | NA |
| *Frasera speciosa* | Plantae | Che-Castaldo; Inouye | Ecosphere | 2011 | 10.1890/ES11-00263.1 |
| *Geum reptans* | Plantae | Weppler; Stoll; Stocklin | J Ecol | 2006 | 10.1111/j.1365-2745.2006.01134.x |
| *Guaiacum sanctum* | Plantae | CITES | Plants Committee | 2008 | NA |
| *Linnaea borealis* | Plantae | Eriksson | J Veg Sci | 1992 | 10.2307/3235999 |
| *Mimulus guttatus* | Plantae | Peterson; Kay; Angert | New Phytol | 2016 | 10.1111/nph.13971 |
| *Paeonia officinalis* | Plantae | Andrieu; Besnard; Fr√©ville; Vaudey; Gauthier; Thompson; Debussche | Biol Conserv | 2017 | 10.1016/j.biocon.2017.08.010 |
| *Panax quinquefolius* | Plantae | Nantel; Gagnon; Nault | Conserv Biol | 1996 | 10.1046/j.1523-1739.1996.10020608.x |
| *Panax quinquefolius* | Plantae | Chandler; McGraw | J Ecol | 2017 | 10.1111/1365-2745.12695 |
| *Psoralea esculenta* | Plantae | Castle | NA | 2006 | NA |
| *Prosartes lanuginosa* | Plantae | Jackson; Pearson; Turner | Forest Ecol Manag | 2013 | 10.1016/j.foreco.2013.05.049 |
| *Rutidosis leptorrhynchoides* | Plantae | Young; Brown; Murray; Thrall; Miller | NA | 2000 | NA |
| *Spathoglottis plicata* | Plantae | Falc√≥n; Ackerman; Tremblay | Biol Invasions | 2017 | 10.1007/s10530-016-1318-8 |
| *Trillium persistens* | Plantae | Plank | NA | 2010 | NA |
| *Vella pseudocytisus subsp. paui* | Plantae | Dominguez Lozano; Moreno Saiz; Schwartz | J Nat Conserv | 2011 | 10.1016/j.jnc.2011.05.005 |
| *Andropogon gerardii* | Plantae | Ott; Hartnett | Am Midl Nat | 2015 | 10.1674/0003-0031-174.1.14 |
| *Anthyllis vulneraria* | Plantae | Davison; Jacquemyn; Adriaens; Honnay; de Kroon; Tuljapurkar | J Ecol | 2010 | 10.1111/j.1365-2745.2009.01611.x |
| *Astragalus scaphoides* | Plantae | Tenhumberg; Crone; Ramula; Tyre | Ecology | 2018 | 10.1002/ecy.2163 |
| *Aurinia saxatilis subsp. saxatilis* | Plantae | ≈†im√°kov√° | NA | 2018 | NA |
| *Bencomia exstipulata* | Plantae | Marrero; Oostermeijer; Nogales; Van Hengstum; Saro; Carqu√©; Sosa; Ba√±ares | J Nat Conserv | 2019 | 10.1016/j.jnc.2018.11.003 |
| *Bromus tectorum* | Plantae | Prev√©y; Seastedt | Popul Ecol | 2015 | 10.1007/s00442-015-3398-z |
| *Centaurea corymbosa* | Plantae | Fr√©ville; Colas; Riba; Caswell; Mignot; Imbert; Olivieri | Ecology | 2004 | 10.1890/03-0119 |
| *Centaurea corymbosa* | Plantae | Belaid; Maurice; Fr√©ville; Carbonell; Imbert | Biol Conserv | 2018 | 10.1016/j.biocon.2018.04.019 |
| *Cephalanthera longifolia* | Plantae | Shefferson; Kull; Tali; Kellett | Ecosphere | 2012 | 10.1890/ES11-00328.1 |
| *Cirsium vulgare* | Plantae | Eckberg; Tenhumberg; Louda | Oecologia | 2014 | 10.1007/s00442-013-2876-4 |
| *Conradina glabra* | Plantae | Bladow | Master Thesis | 2010 | NA |
| *Cypripedium calceolus* | Plantae | Shefferson; Kull; Tali; Kellett | Ecosphere | 2012 | 10.1890/ES11-00328.1 |
| *Cystoseira zosteroides* | Plantae | Capdevila | NA | 2017 | http://hdl.handle.net/2445/117728 |
| *Eritrichium caucasicum* | Plantae | Logofet; Kazantseva; Belova; Onipchenko | Biology Bulletin Reviews | 2018 | 10.1134/S2079086418030076 |
| *Euphorbia telephioides* | Plantae | Ramirez-Bullon | NA | 2016 | NA |
| *Froelichia floridana* | Plantae | McCauley; Ungar | Restor Ecol | 2002 | NA |
| *Gentiana pneumonanthe* | Plantae | Oostermeijer; Brugman; de Boer; den Nijs | J Ecol | 1996 | 10.2307/2261351 |
| *Lepanthes caritensis* | Plantae | Crain; Tremblay; Ferguson | Popul Ecol | 2018 | 10.1002/1438-390X.1002 |
| *Lepanthes rubripetala* | Plantae | Sch√∂delbauerov√°; Tremblay; Kindlmann | Biodivers Conserv | 2010 | 10.1007/s10531-009-9724-1 |
| *Lepanthes rubripetala* | Plantae | Tremblay; Ravent√≥s; Ackerman; | Ann Bot-London | 2015 | 10.1093/aob/mcv031 |
| *Ligularia sibirica* | Plantae | Heinken-Smidova; M√ºnzbergov√° | Folia Geobot | 2012 | 10.1007/s12224-011-9116-7 |
| *Melocactus ernestii* | Plantae | Hughes; Figueira; Jacobi; Borba | Braz J Bot | 2018 | 10.1007/s40415-018-0483-7 |
| *Oncidium poikilostalix* | Plantae | Garc√≠a-Gonz√°lez; Damon; Ravent√≥s; River√≥n-Gir√≥; M√∫jica; Sol√≠s-Montero | Plant Ecol Divers | 2017 | 10.1080/17550874.2017.1315840 |
| *Piriqueta cistoides subsp. caroliniana* | Plantae | Feldman; Morris | J Ecol | 2011 | 10.1111/j.1365-2745.2011.01855.x |
| *Prosartes lanuginosa* | Plantae | Jackson; Pearson; Turner | Forest Ecol Manag | 2013 | 10.1016/j.foreco.2013.05.049 |
| *Serapias cordigera* | Plantae | Pellegrino; Bellusci | Bot J Linn Soc | 2014 | 10.1111/boj.12204 |
| *Silene Ciliata* | Plantae | Gim√©nez-Benavides; Albert; Iriondo; Escudero | Ecography | 2011 | 10.1111/j.1600-0587.2010.06250.x |
| *Trillium grandiflorum* | Plantae | Knight | Am J Bot | 2003 | 10.3732/ajb.90.8.1207 |
| *Tsuga canadensis* | Plantae | Lamar; McGraw | Forest Ecol Manag | 2005 | 10.1016/j.foreco.2005.02.056 |
| *Alaria nana* | Chromalveolata | Pfister; Wang | Ecology | 2005 | 10.1890/04-1952 |
| *Ecklonia radiata* | Chromalveolata | Lees | NA | 2001 | NA |
| *Fucus vesiculosus* | Chromalveolata | Ang; de Wreede | Mar Ecol Prog Ser | 1993 | 10.3354/meps093253 |
| *Aeschynomene virginica* | Plantae | Griffith; Forseth | Ecol Appl | 2005 | 10.1890/02-5219 |
| *Collinsia verna* | Plantae | Kalisz; McPeek | Ecology | 1992 | 10.2307/1940182 |
| *Gilia tenuiflora subsp. hoffmannii* | Plantae | Levine; McEachern; Cowan | J Ecol | 2008 | 10.1111/j.1365-2745.2008.01375.x |
| *Lactuca serriola* | Plantae | Prev√©y; Germino; Huntly | Ecol Appl | 2010 | 10.1890/09-0750 |
| *Malacothrix indecora* | Plantae | Levine; McEachern; Cowan | J Ecol | 2008 | 10.1111/j.1365-2745.2008.01375.x |
| *Phacelia insularis* | Plantae | Levine; McEachern; Cowan | J Ecol | 2008 | 10.1111/j.1365-2745.2008.01375.x |
| *Catopsis compacta* | Plantae | del Castillo; Trujillo-Argueta; Rivera-Garcia; G√≥mez-Ocampo; Mondrag√≥n-Chaparro | Ecol Evol | 2013 | 10.1002/ece3.765 |
| *Catopsis sessiliflora* | Plantae | Winkler; H√ºlber; Hietz | Basic Appl Ecol | 2007 | 10.1016/j.baae.2006.05.003 |
| *Guarianthe aurantiaca* | Plantae | Mondrag√≥n | Plant Spec Biol | 2009 | 10.1111/j.1442-1984.2009.00230.x |
| *Jacquiniella leucomelana* | Plantae | Winkler; H√ºlber; Hietz | Ann Bot-London | 2009 | 10.1093/aob/mcp188 |
| *Jacquiniella teretifolia* | Plantae | Winkler; H√ºlber; Hietz | Ann Bot-London | 2009 | 10.1093/aob/mcp188 |
| *Lepanthes eltoroensis* | Plantae | Tremblay; Ackerman | Biol J Linn Soc | 2001 | 10.1006/bijl.2000.0485 |
| *Lepanthes rubripetala* | Plantae | Tremblay; Ackerman | Biol J Linn Soc | 2001 | 10.1006/bijl.2000.0485 |
| *Lycaste aromatica* | Plantae | Winkler; H√ºlber; Hietz | Ann Bot-London | 2009 | 10.1093/aob/mcp188 |
| *Tillandsia brachycaulos* | Plantae | Mondrag√≥n; Dur√°n; Ram√≠rez; Valverde | J Trop Ecol | 2004 | 10.1017/S0266467403001287 |
| *Tillandsia deppeana* | Plantae | Winkler; H√ºlber; Hietz | Basic Appl Ecol | 2007 | 10.1016/j.baae.2006.05.003 |
| *Tillandsia juncea* | Plantae | Winkler; H√ºlber; Hietz | Basic Appl Ecol | 2007 | 10.1016/j.baae.2006.05.003 |
| *Tillandsia macdougallii* | Plantae | Mondrag√≥n; Ticktin | Conserv Biol | 2011 | 10.1111/j.1523-1739.2011.01691.x |
| *Tillandsia multicaulis* | Plantae | Winkler; H√ºlber; Hietz | Basic Appl Ecol | 2007 | 10.1016/j.baae.2006.05.003 |
| *Tillandsia punctulata* | Plantae | Toledo-Aceves; Valverde; Hern√°ndez-Apolinar | Acta Oecol | 2014 | 10.1016/j.actao.2014.05.009 |
| *Tillandsia recurvata* | Plantae | Valverde; Bernal | Bol Soc Bot Mex | 2010 | 0366-2128 |
| *Tillandsia violacea* | Plantae | Mondrag√≥n; Ticktin | Conserv Biol | 2011 | 10.1111/j.1523-1739.2011.01691.x |
| *Vulpicida pinastri* | Fungi | Shriver; Cutler; Doak | Oecologia | 2012 | 10.1007/s00442-012-2301-4 |
| *Vriesea sanguinolenta* | Plantae | Zotz | Acta Oecol | 2005 | 10.1016/j.actao.2005.05.009 |
| *Asplenium adulterinum* | Plantae | Bucharov√°; M√ºnzbergov√°; T√°jek | Am J Bot | 2010 | 10.3732/ajb.0900351 |
| *Asplenium cuneifolium* | Plantae | Bucharov√°; M√ºnzbergov√°; T√°jek | Am J Bot | 2010 | 10.3732/ajb.0900351 |
| *Asplenium scolopendrium* | Plantae | Bremer; Jongejans | Popul Ecol | 2010 | 10.1007/s10144-009-0143-7 |
| *Actaea elata* | Plantae | Mayberry; Elle | Oecologia | 2010 | 10.1007/s00442-010-1809-8 |
| *Actaea spicata* | Plantae | Fr√∂borg; Eriksson | Can J Bot | 2003 | 10.1139/B03-099 |
| *Adenocarpus gibbsianus* | Plantae | Iriondo; Gim√©nez-Benavides; Albert; Lozano; Escudero | NA | 2009 | 978-84-8014-746-0 |
| *Agrimonia eupatoria* | Plantae | Kiviniemi | Plant Ecol | 2002 | 10.1111/j.1523-1739.2011.01691.x |
| *Alliaria petiolata* | Plantae | Evans; Davis; Raghu; Ragavendran; Landis; Schemske | Ecol Appl | 2012 | 10.1890/11-1291.1 |
| *Anarrhinum fruticosum* | Plantae | Iriondo; Gim√©nez-Benavides; Albert; Lozano; Escudero | NA | 2009 | 978-84-8014-746-0 |
| *Vitaliana primuliflora* | Plantae | Iriondo; Gim√©nez-Benavides; Albert; Lozano; Escudero | NA | 2009 | 978-84-8014-746-0 |
| *Anthericum ramosum* | Plantae | ƒåern√°; M√ºnzbergov√° | PLOS ONE | 2013 | 10.1371/journal.pone.0075563 |
| *Anthyllis vulneraria* | Plantae | Bastrenta; Lebreton; Thompson | J Ecol | 1995 | 10.2307/2261628 |
| *Anthyllis vulneraria* | Plantae | Marcante; Winkler; Erschbamer | Annals Bot | 2009 | 10.1093/aob/mcp047 |
| *Antirrhinum lopesianum* | Plantae | Iriondo; Gim√©nez-Benavides; Albert; Lozano; Escudero | NA | 2009 | 978-84-8014-746-0 |
| *Antirrhinum subbaeticum* | Plantae | Iriondo; Gim√©nez-Benavides; Albert; Lozano; Escudero | NA | 2009 | 978-84-8014-746-0 |
| *Boechera fecunda* | Plantae | Lesica; Shelly | Am J Bot | 1995 | 10.2307/2445615 |
| *Arenaria grandiflora subsp. bolosii* | Plantae | Iriondo; Gim√©nez-Benavides; Albert; Lozano; Escudero | NA | 2009 | 978-84-8014-746-0 |
| *Armeria merinoi* | Plantae | Iriondo; Gim√©nez-Benavides; Albert; Lozano; Escudero | NA | 2009 | 978-84-8014-746-0 |
| *Artemisia genipi* | Plantae | Marcante; Winkler; Erschbamer | Annals Bot | 2009 | 10.1093/aob/mcp047 |
| *Asarum canadense* | Plantae | Damman; Cain | J Ecol | 1998 | 10.1046/j.1365-2745.1998.00242.x |
| *Astragalus peckii* | Plantae | Martin; Meinke | Popul Ecol | 2012 | 10.1007/s10144-012-0318-5 |
| *Astragalus scaphoides* | Plantae | Lesica | Great Basin Nat | 1995 | NA |
| *Astragalus scaphoides* | Plantae | Crone; Lesica | Ecology | 2004 | 10.1890/03-0256 |
| *Astragalus tremolsianus* | Plantae | Iriondo; Gim√©nez-Benavides; Albert; Lozano; Escudero | NA | 2009 | 978-84-8014-746-0 |
| *Astragalus tyghensis* | Plantae | Kaye; Pyke | Ecology | 2003 | 10.1890/0012-9658(2003)084[1464:TEOSTO]2.0.CO;2 |
| *Boltonia decurrens* | Plantae | Smith; Caswell; Mettler-Cherry | Ecol Appl | 2005 | 10.1890/04-0434 |
| *Brassica insularis* | Plantae | Noel; Maurice; Mignot; Gl√©min; Carbonell; Justy; Guyot; Olivieri; Petit | Conserv Genet | 2010 | 10.1007/s10592-010-0056-1 |
| *Calathea ovandensis* | Plantae | Horvitz; Schemske | Ecol Monogr | 1995 | 10.2307/2937136 |
| *Calochortus lyallii* | Plantae | Miller; Antos; Allen | NA | 2004 | NA |
| *Calochortus lyallii* | Plantae | Miller; Antos; Allen | NA | 2004 | NA |
| *Calochortus macrocarpus* | Plantae | Miller; Antos; Allen | NA | 2004 | NA |
| *Carduus nutans* | Plantae | Jongejans; Sheppard; Shea | J Appl Ecol | 2006 | 10.1111/j.1365-2664.2006.01228.x |
| *Carex bigelowii* | Plantae | Carlsson; Callaghan | Oikos | 1991 | 10.2307/3544870 |
| *Carlina vulgaris* | Plantae | Lofgren; Eriksson; Lehtil√§ | Ann Bot Fenn | 2000 | NA |
| *Carlina vulgaris* | Plantae | Jongejans; Jorritsma-Wienk; Becker; Dost√°l; Mild√©n | J Ecol | 2010 | 10.1111/j.1365-2745.2009.01612.x |
| *Centaurea horrida* | Plantae | Pisanu; Farris; Filigheddu; Begona Garcia | Plant Ecol | 2012 | 10.1007/s11258-012-0110-9 |
| *Centaurea jacea* | Plantae | Jongejans; de Kroon | J Ecol | 2005 | 10.1111/j.1365-2745.2005.01003.x |
| *Chamaecrista lineata var. keyensis* | Plantae | Liu; Menges; Quintana-Ascencio | Ecol Appl | 2005 | 10.1890/03-5382 |
| *Chamaelirium luteum* | Plantae | Meagher; Antonovics | Ecology | 1982 | 10.2307/1940111 |
| *Cheirolophus metlesicsii* | Plantae | Iriondo; Gim√©nez-Benavides; Albert; Lozano; Escudero | NA | 2009 | 978-84-8014-746-0 |
| *Actaea elata* | Plantae | Kaye; Pyke | Ecology | 2003 | 10.1890/0012-9658(2003)084[1464:TEOSTO]2.0.CO;2 |
| *Actaea cordifolia* | Plantae | Cook | NA | 1993 | NA |
| *Cirsium dissectum* | Plantae | Jongejans; de Vere; de Kroon | Plant Ecol | 2008 | 10.1007/s11258-008-9397-y |
| *Cirsium pitcheri* | Plantae | Bell; Bowles; McEachern | NA | 2003 | 978-3-642-07869-9 |
| *Cirsium pitcheri* | Plantae | Ellis; Williams; Lesica; Bell; Bierzychudek; Bowles; Crone; Doak; Ehrl√©n; Ellis-Adam; McEachern; Ganesan; Latham; Luijten; Kaye; Knight; Menges; Morris; den Nijs; Oostermeijer; Quintana-Ascencio; Shelly; Stanley; Thorpe; Ticktin; Valverde; Weekley | Ecology | 2012 | 10.1890/11-1052.1 |
| *Cirsium pitcheri* | Plantae | Bell; Powell; Bowles | J Wildlife Manage | 2013 | 10.1002/jwmg.525 |
| *Cirsium pitcheri* | Plantae | Jolls; Marik; Hamz√©; Havens | Biol Conserv | 2015 | 10.1016/j.biocon.2015.04.006 |
| *Cirsium undulatum* | Plantae | Dalgleish; Koons; Adler | J Ecol | 2010 | 10.1111/j.1365-2745.2009.01585.x |
| *Cirsium vulgare* | Plantae | Bullock; Hill; Silvertown | J Ecol | 1994 | 10.2307/2261390 |
| *Cleistesiopsis bifaria* | Plantae | Wells; Willems | NA | 1991 | 90-5103-068-1 |
| *Cleistesiopsis divaricata* | Plantae | Wells; Willems | NA | 1991 | 90-5103-068-1 |
| *Colchicum autumnale* | Plantae | Winter; Jung; Eckstein; Otte; Donath; Kriechbaum | J Appl Ecol | 2014 | 10.1111/1365-2664.12217 |
| *Colchicum autumnale* | Plantae | Winter; Jung; Eckstein; Otte; Donath; Kriechbaum | J Appl Ecol | 2014 | 10.1111/1365-2664.12217 |
| *Corallorhiza trifida* | Plantae | Iriondo; Gim√©nez-Benavides; Albert; Lozano; Escudero | NA | 2009 | 978-84-8014-746-0 |
| *Cryptantha flava* | Plantae | Lucas; Forseth; Casper | J Ecol | 2008 | 10.1111/j.1365-2745.2007.01350.x |
| *Cypripedium calceolus* | Plantae | Garc√≠a; Go√±i; Guzman | Conserv Biol | 2010 | 10.1111/j.1523-1739.2010.01466.x |
| *Cypripedium calceolus* | Plantae | Garc√≠a; Go√±i; Guzman | Conserv Biol | 2010 | 10.1111/j.1523-1739.2010.01466.x |
| *Cypripedium fasciculatum* | Plantae | Thorpe; Stanley; Kayne; Latham | NA | 2011 | NA |
| *Danthonia sericea* | Plantae | Moloney | Ecology | 1988 | 10.2307/1941656 |
| *Dicentra canadensis* | Plantae | Lin; Miriti; Goodell | Ecol Evol | 2016 | 10.1002/ece3.2163 |
| *Dicerandra frutescens* | Plantae | Menges; Quintana-Ascencio; Weekley; Gaoue | Biol Conserv | 2006 | 10.1016/j.biocon.2005.08.002 |
| *Dipsacus fullonum* | Plantae | Werner; Caswell | Ecology | 1977 | 10.2307/1936930 |
| *Dorycnium spectabile* | Plantae | Iriondo; Gim√©nez-Benavides; Albert; Lozano; Escudero | NA | 2009 | 978-84-8014-746-0 |
| *Draba asterophora* | Plantae | Putnam | NA | 2013 | NA |
| *Dracocephalum austriacum* | Plantae | Andrello | NA | 2010 | NA |
| *Echinacea angustifolia* | Plantae | Dalgleish; Koons; Adler | J Ecol | 2010 | 10.1111/j.1365-2745.2009.01585.x |
| *Echinacea angustifolia* | Plantae | Hurlburt | NA | 1999 | NA |
| *Echinospartum ibericum subsp. algibicum* | Plantae | Iriondo; Gim√©nez-Benavides; Albert; Lozano; Escudero | NA | 2009 | 978-84-8014-746-0 |
| *Eriogonum longifolium var. gnaphalifolium* | Plantae | Satterthwaite; Menges; Quintana-Ascencio | Ecol Appl | 2002 | 10.1890/1051-0761(2002)012[1672:ASBPVI]2.0.CO;2 |
| *Erodium paularense* | Plantae | Iriondo; Gim√©nez-Benavides; Albert; Lozano; Escudero | NA | 2009 | 978-84-8014-746-0 |
| *Eryngium alpinum* | Plantae | Andrello; Bizoux; Barbet-Massin; Gaudeul; Nicol√®; Till-Bottraud | Biol Conserv | 2012 | 10.1016/j.biocon.2011.12.012 |
| *Eryngium cuneifolium* | Plantae | Menges; Quintana-Ascencio | Ecol Monogr | 2004 | 10.1890/03-4029 |
| *Eupatorium perfoliatum* | Plantae | Byers; Meagher | Ecol Appl | 1997 | 10.1890/1051-0761(1997)007[0519:ACODCI]2.0.CO;2 |
| *Eupatorium resinosum* | Plantae | Byers; Meagher | Ecol Appl | 1997 | 10.1890/1051-0761(1997)007[0519:ACODCI]2.0.CO;2 |
| *Oenothera coloradensis subsp. coloradensis* | Plantae | Floyd; Ranker | Int J Plant Sci | 1998 | 10.1086/297607 |
| *Gentiana pneumonanthe* | Plantae | Oostermeijer; Brugman; de Boer; den Nijs | J Ecol | 1996 | 10.2307/2261351 |
| *Geranium sylvaticum* | Plantae | Ramula; Toivonen; Mutikainen | Int J Plant Sci | 2007 | 10.1086/512040 |
| *Geum rivale* | Plantae | Kiviniemi | Plant Ecol | 2002 | 10.1111/j.1523-1739.2011.01691.x |
| *Pyrrocoma radiata* | Plantae | Kaye; Pyke | Ecology | 2003 | 10.1890/0012-9658(2003)084[1464:TEOSTO]2.0.CO;2 |
| *Stenaria nigricans* | Plantae | Dalgleish; Koons; Adler | J Ecol | 2010 | 10.1111/j.1365-2745.2009.01585.x |
| *Helianthemum polygonoides* | Plantae | Iriondo; Gim√©nez-Benavides; Albert; Lozano; Escudero | NA | 2009 | 978-84-8014-746-0 |
| *Helianthemum teneriffae* | Plantae | Iriondo; Gim√©nez-Benavides; Albert; Lozano; Escudero | NA | 2009 | 978-84-8014-746-0 |
| *Heliconia acuminata* | Plantae | Bruna | Ecology | 2003 | 10.1890/0012-9658(2003)084[0932:APPIFH]2.0.CO;2 |
| *Heliconia metallica* | Plantae | Schleuning; Huam√°n; Matthies | J Ecol | 2008 | 10.1111/j.1365-2745.2008.01416.x |
| *Hilaria mutica* | Plantae | Vega; Monta√±a | Plant Ecol | 2004 | 10.1023/B:VEGE.0000048094.21994.74 |
| *Horkelia congesta* | Plantae | Kaye; Benfield | NA | 2004 | NA |
| *Hydrastis canadensis* | Plantae | Sinclair | NA | 2002 | NA |
| *Tetraneuris herbacea* | Plantae | Campbell; Husband | Heredity | 2005 | 10.1038/sj.hdy.6800653 |
| *Hyparrhenia diplandra* | Plantae | Garnier; Dajoz | J Ecol | 2001 | 10.1890/0012-9658(2001)082[1720:ESOALV]2.0.CO;2 |
| *Hypericum cumulicola* | Plantae | Quintana-Ascencio; Menges; Weekley | Conserv Biol | 2003 | 10.1046/j.1523-1739.2003.01431.x |
| *Jurinea fontqueri* | Plantae | Iriondo; Gim√©nez-Benavides; Albert; Lozano; Escudero | NA | 2009 | 978-84-8014-746-0 |
| *Kosteletzkya pentacarpos* | Plantae | Pino; Pic√≥; Roa | Bot J Linn Soc | 2007 | 10.1111/j.1095-8339.2007.00628.x |
| *Kunkeliella subsucculenta* | Plantae | Iriondo; Gim√©nez-Benavides; Albert; Lozano; Escudero | NA | 2009 | 978-84-8014-746-0 |
| *Laserpitium longiradium* | Plantae | Iriondo; Gim√©nez-Benavides; Albert; Lozano; Escudero | NA | 2009 | 978-84-8014-746-0 |
| *Lathyrus Vernus* | Plantae | Ehrl√©n | J Ecol | 1995 | 10.2307/2261568 |
| *Leontopodium nivale subsp. alpinum* | Plantae | Keller; Vittoz | Alpine Bot | 2015 | 10.1007/s00035-014-0142-y |
| *Lepanthes rupestris* | Plantae | Tremblay; Ackerman | Biol J Linn Soc | 2001 | 10.1006/bijl.2000.0485 |
| *Lepanthes rupestris* | Plantae | Tremblay; McCarthy | PLOS ONE | 2014 | 10.1371/journal.pone.0102859 |
| *Lepidium davisii* | Plantae | Bernatus | NA | 1995 | NA |
| *Physaria ovalifolia* | Plantae | Dalgleish; Koons; Adler | J Ecol | 2010 | 10.1111/j.1365-2745.2009.01585.x |
| *Liatris scariosa* | Plantae | Ellis | Ecology | 2012 | 10.1890/11-1052.1 |
| *Limonium erectum* | Plantae | Iriondo; Gim√©nez-Benavides; Albert; Lozano; Escudero | NA | 2009 | 978-84-8014-746-0 |
| *Limonium geronense* | Plantae | Iriondo; Gim√©nez-Benavides; Albert; Lozano; Escudero | NA | 2009 | 978-84-8014-746-0 |
| *Limonium malacitanum* | Plantae | Iriondo; Gim√©nez-Benavides; Albert; Lozano; Escudero | NA | 2009 | 978-84-8014-746-0 |
| *Linum flavum* | Plantae | M√ºnzbergov√° | Plant Biology | 2013 | 10.1111/plb.12007 |
| *Linum tenuifolium* | Plantae | M√ºnzbergov√° | Plant Biology | 2013 | 10.1111/plb.12007 |
| *Lithospermum ruderale* | Plantae | Bricker; Maron | Ecology | 2012 | 10.1890/11-0948.1 |
| *Lomatium bradshawii* | Plantae | Kaye; Pyke | Ecology | 2003 | 10.1890/0012-9658(2003)084[1464:TEOSTO]2.0.CO;2 |
| *Lomatium bradshawii* | Plantae | Kaye; Pendergrass; Finley; Kauffman | Ecol Appl | 2001 | 10.1890/1051-0761(2001)011[1366:TEOFOT]2.0.CO;2 |
| *Lomatium cookii* | Plantae | Kaye; Pyke | Ecology | 2003 | 10.1890/0012-9658(2003)084[1464:TEOSTO]2.0.CO;2 |
| *Lotus arinagensis* | Plantae | Iriondo; Gim√©nez-Benavides; Albert; Lozano; Escudero | NA | 2009 | 978-84-8014-746-0 |
| *Lupinus lepidus* | Plantae | Bishop | NA | 1996 | NA |
| *Lupinus tidestromii* | Plantae | Dangremond; Knight | Ecology | 2010 | 10.1890/09-0418.1 |
| *Mimulus cardinalis* | Plantae | Angert | Ecology | 2006 | 10.1890/0012-9658(2006)87[2014:DOCAMP]2.0.CO;2 |
| *Molinia caerulea* | Plantae | Jacquemyn; Brys; Neubert | Ecol Appl | 2005 | 10.1890/04-1762 |
| *Narcissus pseudonarcissus* | Plantae | Barkham | J Ecol | 1980 | 10.2307/2259425 |
| *Neotinea ustulata* | Plantae | Shefferson; Tali | J Ecol | 2007 | 10.1111/j.1365-2745.2006.01195.x |
| *Oenothera deltoides* | Plantae | Thomson | Conserv Biol | 2005 | 10.1111/j.1523-1739.2005.004108.x |
| *Orchis purpurea* | Plantae | Jacquemyns; Brys; Jongejans | Ecology | 2010 | 10.1890/08-2321.1 |
| *Oxalis acetosella* | Plantae | Berg | Ecography | 2002 | 10.1034/j.1600-0587.2002.250211.x |
| *Oxytropis jabalambrensis* | Plantae | Iriondo; Gim√©nez-Benavides; Albert; Lozano; Escudero | NA | 2009 | 978-84-8014-746-0 |
| *Panax quinquefolius* | Plantae | Shahi | NA | 2007 | NA |
| *Panax quinquefolius* | Plantae | Charron; Gagnon | J Ecol | 1991 | 10.2307/2260724 |
| *Parolinia glabriuscula* | Plantae | Iriondo; Gim√©nez-Benavides; Albert; Lozano; Escudero | NA | 2009 | 978-84-8014-746-0 |
| *Paronychia jamesii* | Plantae | Dalgleish; Koons; Adler | J Ecol | 2010 | 10.1111/j.1365-2745.2009.01585.x |
| *Petrocoptis pyrenaica subsp. pseudoviscosa* | Plantae | Garc√≠a; Guzman; Go√±i | Biol Conserv | 2002 | 10.1016/S0006-3207(01)00113-6 |
| *Petrocoptis pyrenaica subsp. pseudoviscosa* | Plantae | Garc√≠a; Guzman; Go√±i | Biol Conserv | 2002 | 10.1016/S0006-3207(01)00113-6 |
| *Pimpinella saxifraga* | Plantae | Auestad; Rydgren; Jongejans; Kroon | Biol Conserv | 2010 | 10.1016/j.biocon.2009.12.037 |
| *Pinguicula ionantha* | Plantae | Kesler; Trusty; Hermann; Guyer | Oecologia | 2008 | 10.1007/s00442-008-1022-1 |
| *Plantago coronopus* | Plantae | Waite | J Ecol | 1984 | 10.2307/2259533 |
| *Plantago coronopus* | Plantae | Villellas; Ehrl√©n; Olesen; Braza; Garc√≠a | Ecography | 2013 | 10.1111/j.1600-0587.2012.07425.x |
| *Plantago coronopus* | Plantae | Villellas; Ehrl√©n; Olesen; Braza; Garc√≠a | Ecography | 2013 | 10.1111/j.1600-0587.2012.07425.x |
| *Plantago coronopus* | Plantae | Villellas; Ehrl√©n; Olesen; Braza; Garc√≠a | Ecography | 2013 | 10.1111/j.1600-0587.2012.07425.x |
| *Plantago coronopus* | Plantae | Villellas; Ehrl√©n; Olesen; Braza; Garc√≠a | Ecography | 2013 | 10.1111/j.1600-0587.2012.07425.x |
| *Plantago coronopus* | Plantae | Villellas; Ehrl√©n; Olesen; Braza; Garc√≠a | Ecography | 2013 | 10.1111/j.1600-0587.2012.07425.x |
| *Plantago media* | Plantae | Eriksson; Eriksson | J Veg Sci | 2000 | 10.2307/3236803 |
| *Poa alpina* | Plantae | Marcante; Winkler; Erschbamer | Annals Bot | 2009 | 10.1093/aob/mcp047 |
| *Polemonium van-bruntiae* | Plantae | Bermingham | Plant Ecol | 2010 | 10.1007/s11258-010-9762-5 |
| *Polygonella basiramia* | Plantae | Maliakal Witt | NA | 2004 | NA |
| *Potentilla anserina* | Plantae | Eriksson | J Ecol | 1988 | 10.2307/2260610 |
| *Primula elatior* | Plantae | Jacquemyn; Brys | Ecology | 2008 | 10.1890/07-1908.1 |
| *Primula farinosa* | Plantae | Lindborg; Ehrl√©n | Conserv Biol | 2002 | 10.1046/j.1523-1739.2002.00509.x |
| *Primula veris* | Plantae | Lehtil√§; Syrj√§nen; Leimu; Garc√≠a; Ehrl√©n | Conserv Biol | 2006 | 10.1111/j.1523-1739.2006.00368.x |
| *Primula veris* | Plantae | Lehtil√§; Syrj√§nen; Leimu; Garc√≠a; Ehrl√©n | Conserv Biol | 2006 | 10.1111/j.1523-1739.2006.00368.x |
| *Primula veris* | Plantae | Endels; Jacquemyn; Brys; Hermy | Plant Ecol | 2005 | 10.1007/s11258-004-0026-0 |
| *Primula vulgaris* | Plantae | Valverde; Silvertown | J Ecol | 1998 | 10.1046/j.1365-2745.1998.00280.x |
| *Primula vulgaris* | Plantae | Valdes; Garc√≠a; Garc√≠a; Ehrl√©n | Ecography | 2013 | 10.1111/j.1600-0587.2013.00216.x |
| *Pseudomisopates rivas-martinezii* | Plantae | Iriondo; Gim√©nez-Benavides; Albert; Lozano; Escudero | NA | 2009 | 978-84-8014-746-0 |
| *Psoralea tenuiflora* | Plantae | Dalgleish; Koons; Adler | J Ecol | 2010 | 10.1111/j.1365-2745.2009.01585.x |
| *Pyrrocoma radiata* | Plantae | Pfingsten | NA | 2013 | NA |
| *Ramonda myconi* | Plantae | Pic√≥; Riba | Plant Ecol | 2002 | 10.1023/A:1020310609348 |
| *Ranunculus peltatus* | Plantae | Idestam-Almquist | NA | 1998 | NA |
| *Ratibida columnifera* | Plantae | Dalgleish; Koons; Adler | J Ecol | 2010 | 10.1111/j.1365-2745.2009.01585.x |
| *Rubus praecox* | Plantae | Lambrecht-McDowell; Radosevich | Biol Invasions | 2005 | 10.1007/s10530-004-0870-9 |
| *Rumex rupestris* | Plantae | Iriondo; Gim√©nez-Benavides; Albert; Lozano; Escudero | NA | 2009 | 978-84-8014-746-0 |
| *Santolina melidensis* | Plantae | Iriondo; Gim√©nez-Benavides; Albert; Lozano; Escudero | NA | 2009 | 978-84-8014-746-0 |
| *Saponaria bellidifolia* | Plantae | Cserg≈ë; Moln√°r; Garc√≠a | Popul Ecol | 2011 | 10.1007/s10144-010-0249-y |
| *Sarcocapnos baetica* | Plantae | Salinas; Su√°rez; Blanca | Can J Bot | 2002 | 10.1139/b02-013 |
| *Sarcocapnos enneaphylla* | Plantae | Salinas; Su√°rez; Blanca | Can J Bot | 2002 | 10.1139/b02-013 |
| *Sarcocapnos pulcherrima* | Plantae | Salinas; Su√°rez; Blanca | Can J Bot | 2002 | 10.1139/b02-013 |
| *Sarracenia purpurea* | Plantae | Gotelli; Ellison | Ecol Appl | 2006 | 10.1890/04-0479 |
| *Saussurea medusa* | Plantae | Law; Salick; Knight | Plant Ecol | 2010 | 10.1007/s11258-010-9761-6 |
| *Saxifraga aizoides* | Plantae | Marcante; Winkler; Erschbamer | Annals Bot | 2009 | 10.1093/aob/mcp047 |
| *Saxifraga cotyledon* | Plantae | Dinnetz; Nilsson | Plant Ecol | 2002 | 10.1023/A:1015593311183 |
| *Silene douglasii var. oraria* | Plantae | Kephart; Paladino | Am J Bot | 1997 | 10.2307/2446079 |
| *Solidago mollis* | Plantae | Dalgleish; Koons; Adler | J Ecol | 2010 | 10.1111/j.1365-2745.2009.01585.x |
| *Sphaeralcea coccinea* | Plantae | Dalgleish; Koons; Adler | J Ecol | 2010 | 10.1111/j.1365-2745.2009.01585.x |
| *Sporobolus heterolepis* | Plantae | Dalgleish; Kula; Hartnett; Sandercock | Am J Bot | 2008 | 10.3732/ajb.2007277 |
| *Succisa pratensis* | Plantae | Wallin; Svensson | Folia Geobot | 2012 | 10.1007/s12224-012-9123-3 |
| *Succisa pratensis* | Plantae | Mild√©n | NA | 2005 | NA |
| *Taraxacum campylodes* | Plantae | Vavrek; McGraw; Yang | J Ecol | 1997 | 10.2307/2960501 |
| *Thelesperma megapotamicum* | Plantae | Dalgleish; Koons; Adler | J Ecol | 2010 | 10.1111/j.1365-2745.2009.01585.x |
| *Thymus vulgaris* | Plantae | Iriondo; Gim√©nez-Benavides; Albert; Lozano; Escudero | NA | 2009 | 978-84-8014-746-0 |
| *Trillium grandiflorum* | Plantae | Schmucki | NA | 2009 | NA |
| *Trillium ovatum* | Plantae | Ream | NA | 2011 | NA |
| *Trollius europaeus* | Plantae | Lemke; Salguero-G√≥mez | Popul Ecol | 2015 | 10.1007/s10144-015-0519-9 |
| *Trollius laxus* | Plantae | Scanga; Leopold | Biol Conserv | 2012 | 10.1016/j.biocon.2012.01.061 |
| *Trollius laxus* | Plantae | Scanga | Plant Ecol | 2014 | 10.1007/s11258-014-0344-9 |
| *Verbascum fontqueri* | Plantae | Iriondo; Gim√©nez-Benavides; Albert; Lozano; Escudero | NA | 2009 | 978-84-8014-746-0 |
| *Zea diploperennis* | Plantae | S√°nchez-Vel√°squez; Ezcurra; Martinez-Ramos; Alvarez-Buylla; Lorente | J Ecol | 2002 | 10.1046/j.1365-2745.2002.00702.x |
| *Alyxia stellata* | Plantae | Wong; Ticktin | Environ Conserv | 2014 | 10.1017/S0376892914000204 |
| *Machaerium cuspidatum* | Plantae | Nabe-Nielsen | J Trop Ecol | 2004 | 10.1017/S0266467404001609 |
| *Borassus aethiopum* | Plantae | Barot; Gignoux; Vuattoux | J Trop Ecol | 2000 | 10.1017/S0266467400001620 |
| *Ceratozamia mirandae* | Plantae | P√©rez-Farrera; Vovides; Octavio-Aguilar; Gonz√°lez-Astorga; Cruz-Rodr√≠guez; Hern√°ndez-Jonap√°; Villalobos-M√©ndez | Plant Ecol | 2006 | 10.1007/s11258-006-9135-2 |
| *Chamaedorea elegans* | Plantae | Valverde; Hern√°ndez-Apolinar; Mendoza-Amarom | J Sustain Forest | 2006 | 10.1300/J091v23n01_05 |
| *Chamaedorea radicalis* | Plantae | Endress; Gorchov; Robert; Noble | Ecol Appl | 2004 | 10.1890/02-5365 |
| *Chamaedorea radicalis* | Plantae | Berry; Gorchov; Endress; Stevens | Popul Ecol | 2008 | 10.1007/s10144-007-0067-z |
| *Dioon merolae* | Plantae | L√°zaro-Zerme√±o; Gonz√°lez-Espinosa; Mendoza; Martinez-Ramos; Quintana-Ascencio | Forest Ecol Manag | 2011 | 10.1016/j.foreco.2010.10.028 |
| *Eremospatha macrocarpa* | Plantae | Kouassi; Barot; Gignoux; Bi | J Trop Ecol | 2008 | 10.1017/S0266467408005312 |
| *Euterpe edulis* | Plantae | Silva-Matos; Freckleton; Watkinson | Ecology | 1999 | 10.1890/0012-9658(1999)080[2635:TRODDI]2.0.CO;2 |
| *Euterpe oleracea* | Plantae | Arango; Duque; Mu√±oz | Int J Trop Biol | 2010 | 10.15517/rbt.v58i1.5222 |
| *Geonoma pohliana subsp. weddelliana* | Plantae | Souza; Martins | Austral Ecol | 2006 | 10.1111/j.1442-9993.2006.01650.x |
| *Geonoma schottiana* | Plantae | Sampaio; Scariot | J Trop Ecol | 2010 | 10.1017/S0266467409990599 |
| *Iriartea deltoidea* | Plantae | Pinard | Biotropica | 1993 | 10.2307/2388974 |
| *Laccosperma secundiflorum* | Plantae | Kouassi; Barot; Gignoux; Bi | J Trop Ecol | 2008 | 10.1017/S0266467408005312 |
| *Pseudophoenix sargentii* | Plantae | Dur√°n; Franco | NA | 1992 | NA |
| *Sabal yapa* | Plantae | Pulido; Valverde; Caballero | J Trop Ecol | 2007 | 10.1017/S0266467406003877 |
| *Thrinax radiata* | Plantae | Olmsted; Alvarez-Buylla | Ecol Appl | 1995 | 10.2307/1942038 |
| *Zamia amblyphyllidia* | Plantae | Negron-Ortiz; Gorchov; Breckon | Int J Plant Sci | 1996 | 10.1086/297381 |
| *Acacia suaveolens* | Plantae | Warton; Wardle | Austral Ecol | 2003 | 10.1046/j.1442-9993.2003.01246.x |
| *Ardisia elliptica* | Plantae | Koop; Horvitz | Ecology | 2005 | 10.1890/04-1483 |
| *Atriplex acanthocarpa* | Plantae | Verhulst; Monta√±a; Mandujano; Franco | Oecologia | 2008 | 10.1007/s00442-008-0980-7 |
| *Atriplex canescens* | Plantae | Verhulst; Monta√±a; Mandujano; Franco | Oecologia | 2008 | 10.1007/s00442-008-0980-7 |
| *Clidemia hirta* | Plantae | DeWalt | Biol Invasions | 2006 | 10.1007/s10530-005-5277-8 |
| *Cytisus scoparius* | Plantae | Neubert; Parker | Risk Anal | 2004 | 10.1111/j.0272-4332.2004.00481.x |
| *Fumana procumbens* | Plantae | Bengtsson | J Ecol | 1993 | 10.2307/2261672 |
| *Gardenia actinocarpa* | Plantae | Osunkoya | Biol Conserv | 2003 | 10.1016/S0006-3207(02)00417-2 |
| *Helianthemum juliae* | Plantae | Marrero-G√≥mez; Oostermeijer; Carqu√©-√Ålamo; Ba√±ares-Baudet | Biol Conserv | 2007 | 10.1016/j.biocon.2007.01.010 |
| *Persoonia bargoensis* | Plantae | McKenna | NA | 2007 | NA |
| *Persoonia glaucescens* | Plantae | McKenna | NA | 2007 | NA |
| *Purshia subintegra* | Plantae | Maschinski; Baggs; Quintana-Ascencio; Menges | Conserv Biol | 2006 | 10.1111/j.1523-1739.2006.00272.x |
| *Rosmarinus tomentosus* | Plantae | Iriondo; Gim√©nez-Benavides; Albert; Lozano; Escudero | NA | 2009 | 978-84-8014-746-0 |
| *Tetramolopium arenarium* | Plantae | Aplet; Laven; Shaw | Nat Area J | 1994 | NA |
| *Vella pseudocytisus subsp. paui* | Plantae | Iriondo; Gim√©nez-Benavides; Albert; Lozano; Escudero | NA | 2009 | 978-84-8014-746-0 |
| *Vella pseudocytisus subsp. paui* | Plantae | Iriondo; Gim√©nez-Benavides; Albert; Lozano; Escudero | NA | 2009 | 978-84-8014-746-0 |
| *Astrophytum asterias* | Plantae | Martinez-Avalos | NA | 2007 | NA |
| *Astrophytum ornatum* | Plantae | Zepeda-Martinez; Mandujano; Mandujano; Golubov | J Arid Environ | 2013 | 10.1016/j.jaridenv.2012.08.006 |
| *Escobaria robbinsorum* | Plantae | Schmalzel; Reichenbacher; Rutman | Madrono | 1995 | NA |
| *Escontria chiotilla* | Plantae | Ortega-Baes | NA | 2001 | NA |
| *Euphorbia fontqueriana* | Plantae | Iriondo; Gim√©nez-Benavides; Albert; Lozano; Escudero | NA | 2009 | 978-84-8014-746-0 |
| *Mammillaria crucigera* | Plantae | Contreras; Valverde | J Arid Environ | 2002 | 10.1006/jare.2001.0926 |
| *Mammillaria gaumeri* | Plantae | Ferrer; Dur√°n; M√©ndez; Dorantes; Dzib | Bol Soc Bot Mex | 2011 | NA |
| *Mammillaria hernandezii* | Plantae | Rodriguez Ortega | NA | 2008 | NA |
| *Mammillaria huitzilopochtli* | Plantae | Flores Mart√≠nez; Manzanero-Medino; Golubov; Monta√±a; Mandujano | Plant Ecol | 2010 | 10.1007/s11258-010-9737-6 |
| *Mammillaria huitzilopochtli* | Plantae | Flores Mart√≠nez | NA | 2010 | NA |
| *Mammillaria magnimamma* | Plantae | Valverde; Quijas; Lopez-Villavicencio; Castillo | Plant Ecol | 2004 | 10.1023/B:VEGE.0000021662.78634.de |
| *Mammillaria napina* | Plantae | Rodriguez Ortega | NA | 2008 | NA |
| *Mammillaria solisioides* | Plantae | Rodriguez Ortega | NA | 2008 | NA |
| *Neobuxbaumia macrocephala* | Plantae | Esparza-Olgu√≠n; Valverde; Mandujano | Popul Ecol | 2005 | 10.1007/s10144-005-0230-3 |
| *Neobuxbaumia macrocephala* | Plantae | Esparza Olgu√≠n | NA | 2005 | NA |
| *Neobuxbaumia mezcalaensis* | Plantae | Esparza-Olgu√≠n; Valverde; Mandujano | Popul Ecol | 2005 | 10.1007/s10144-005-0230-3 |
| *Neobuxbaumia mezcalaensis* | Plantae | Esparza Olgu√≠n | NA | 2005 | NA |
| *Neobuxbaumia polylopha* | Plantae | Arroyo-Cosultchi; Golubov; Mandujano | Acta Oecol | 2016 | 10.1016/j.actao.2016.01.006 |
| *Neobuxbaumia tetetzo* | Plantae | Esparza-Olgu√≠n; Valverde; Mandujano | Popul Ecol | 2005 | 10.1007/s10144-005-0230-3 |
| *Neobuxbaumia tetetzo* | Plantae | Esparza Olgu√≠n | NA | 2005 | NA |
| *Opuntia macrorhiza* | Plantae | Haridas; Keeler; Tenhumberg | Ecology | 2015 | 10.1890/13-1984.1 |
| *Opuntia rastrera* | Plantae | Mandujano; Monta√±a; Franco; Golubov; Flores-Mart√≠nez | Ecology | 2001 | 10.2307/2679864 |
| *Pediocactus bradyi* | Plantae | Shryock; Esque; Hughes | Am J Bot | 2014 | 10.3732/ajb.1400035 |
| *Pterocereus gaumeri* | Plantae | M√©ndez; Dur√°n; Olmsted | Biotropica | 2004 | 10.1646/1601 |
| *Stenocereus eruca* | Plantae | Clark-Tapia | NA | 2004 | NA |
| *Acer saccharum* | Plantae | Lin; Augspurger | Forest Ecol Manag | 2008 | 10.1016/j.foreco.2008.02.040 |
| *Aesculus turbinata* | Plantae | Kaneko; Takada; Kawano | Plant Spec Biol | 1999 | 10.1046/j.1442-1984.1999.00007.x |
| *Alnus incana subsp. rugosa* | Plantae | Huenneke; Marks | Ecology | 1987 | 10.2307/1939207 |
| *Bursera glabrifolia* | Plantae | Hern√°ndez-Apolinar; Valverde; Purata | Forest Ecol Manag | 2006 | 10.1016/j.foreco.2005.10.072 |
| *Castanea dentata* | Plantae | Davelos; Jarosz | J Ecol | 2004 | 10.1111/j.0022-0477.2004.00907.x |
| *Fagus grandifolia* | Plantae | da Silva Batista; Platt; Macchiavelli | Ecology | 1998 | 10.2307/176863 |
| *Khaya senegalensis* | Plantae | Gaoue; Ticktin | Conserv Biol | 2010 | 10.1111/j.1523-1739.2009.01345.x |
| *Magnolia macrophylla var. dealbata* | Plantae | S√°nchez-Vel√°squez; Pineda-L√≥pez | Popul Ecol | 2010 | 10.1007/s10144-009-0161-5 |
| *Manilkara zapota* | Plantae | Cruz-Rodr√≠guez; Lopez-Villavicencio; Valverde | J Trop Ecol | 2009 | 10.1017/S0266467408005713 |
| *Phyllanthus emblica* | Plantae | Ticktin; Ganesan; Paramesha; Setty | J Appl Ecol | 2012 | 10.1111/j.1365-2664.2012.02156.x |
| *Phyllanthus emblica* | Plantae | Ellis; Williams; Lesica; Bell; Bierzychudek; Bowles; Crone; Doak; Ehrl√©n; Ellis-Adam; McEachern; Ganesan; Latham; Luijten; Kaye; Knight; Menges; Morris; den Nijs; Oostermeijer; Quintana-Ascencio; Shelly; Stanley; Thorpe; Ticktin; Valverde; Weekley | Ecology | 2012 | 10.1890/11-1052.1 |
| *Phyllanthus indofischeri* | Plantae | Ticktin; Ganesan; Paramesha; Setty | J Appl Ecol | 2012 | 10.1111/j.1365-2664.2012.02156.x |
| *Pinus lambertiana* | Plantae | van Mantgem; Stephenson | J Ecol | 2005 | 10.1111/j.1365-2745.2005.01007.x |
| *Pinus lambertiana* | Plantae | Maloney; Vogler; Eckert; Jensen; Neale | Forest Ecol Manag | 2011 | 10.1016/j.foreco.2011.05.011 |
| *Pinus nigra* | Plantae | Buckley; Brockerhoff; Langer; Ledgard; North; Rees | J Appl Ecol | 2005 | 10.1111/j.1365-2664.2005.01100.x |
| *Pinus strobus* | Plantae | M√ºnzbergov√°; Hadincov√°; Wild; Kindlmannov√° | PLOS ONE | 2013 | 10.1371/journal.pone.0056953 |
| *Prioria copaifera* | Plantae | Condit | Forest Ecol Manag | 1993 | 10.1016/0378-1127(93)90045-O |
| *Prosopis laevigata* | Plantae | Bernal | NA | 2004 | NA |
| *Prunus africana* | Plantae | Stewart | NA | 2001 | NA |
| *Rhododendron ponticum* | Plantae | Salguero-G√≥mez | NA | 2004 | NA |
| *Rhododendron ponticum* | Plantae | Salguero-G√≥mez | NA | 2004 | NA |
| *Sapium sebiferum* | Plantae | Renne | NA | 2001 | NA |
| *Shorea leprosula* | Plantae | Chen; Visser; Jongejans; van Breugel; Zuidema; Kassim; de Kroon | J Ecol | 2011 | 10.1111/j.1365-2745.2011.01825.x |
| *Styrax obassis* | Plantae | Abe; Nokashizuka; Tanoka | J Veg Sci | 1998 | 10.2307/3237044 |
| *Taxus floridana* | Plantae | Kwit; Horvitz; Platt | Conserv Biol | 2004 | 10.1111/j.1523-1739.2004.00567.x |
| *Tsuga canadensis* | Plantae | Lamar; McGraw | Forest Ecol Manag | 2005 | 10.1016/j.foreco.2005.02.056 |
| *Ziziphus jujuba* | Plantae | Zull; Lawes; Cacho | Environ Modell Softw | 2015 | 10.1016/j.envsoft.2015.10.026 |
| *Oeceoclades maculata* | Plantae | River√≥n-Gir√≥; Ravent√≥s; Damon; Garc√≠a-Gonz√°lez; M√∫jica | Biol Invasions | 2019 | 10.1007/s10530-019-01945-7 |
| *Malacothrix indecora* | Plantae | Levine; McEachern; Cowan | J Ecol | 2008 | 10.1111/j.1365-2745.2008.01375.x |
| *Phacelia insularis* | Plantae | Levine; McEachern; Cowan | J Ecol | 2008 | 10.1111/j.1365-2745.2008.01375.x |
| *Catopsis compacta* | Plantae | del Castillo; Trujillo-Argueta; Rivera-Garcia; G√≥mez-Ocampo; Mondrag√≥n-Chaparro | Ecol Evol | 2013 | 10.1002/ece3.765 |
| *Catopsis sessiliflora* | Plantae | Winkler; H√ºlber; Hietz | Basic Appl Ecol | 2007 | 10.1016/j.baae.2006.05.003 |
| *Guarianthe aurantiaca* | Plantae | Mondrag√≥n | Plant Spec Biol | 2009 | 10.1111/j.1442-1984.2009.00230.x |
| *Jacquiniella leucomelana* | Plantae | Winkler; H√ºlber; Hietz | Ann Bot-London | 2009 | 10.1093/aob/mcp188 |
| *Jacquiniella teretifolia* | Plantae | Winkler; H√ºlber; Hietz | Ann Bot-London | 2009 | 10.1093/aob/mcp188 |
| *Lepanthes eltoroensis* | Plantae | Tremblay; Ackerman | Biol J Linn Soc | 2001 | 10.1006/bijl.2000.0485 |
| *Lepanthes rubripetala* | Plantae | Tremblay; Ackerman | Biol J Linn Soc | 2001 | 10.1006/bijl.2000.0485 |
| *Lycaste aromatica* | Plantae | Winkler; H√ºlber; Hietz | Ann Bot-London | 2009 | 10.1093/aob/mcp188 |
| *Tillandsia brachycaulos* | Plantae | Mondrag√≥n; Dur√°n; Ram√≠rez; Valverde | J Trop Ecol | 2004 | 10.1017/S0266467403001287 |
| *Tillandsia deppeana* | Plantae | Winkler; H√ºlber; Hietz | Basic Appl Ecol | 2007 | 10.1016/j.baae.2006.05.003 |
| *Tillandsia juncea* | Plantae | Winkler; H√ºlber; Hietz | Basic Appl Ecol | 2007 | 10.1016/j.baae.2006.05.003 |
| *Tillandsia macdougallii* | Plantae | Mondrag√≥n; Ticktin | Conserv Biol | 2011 | 10.1111/j.1523-1739.2011.01691.x |
| *Tillandsia multicaulis* | Plantae | Winkler; H√ºlber; Hietz | Basic Appl Ecol | 2007 | 10.1016/j.baae.2006.05.003 |
| *Tillandsia punctulata* | Plantae | Toledo-Aceves; Valverde; Hern√°ndez-Apolinar | Acta Oecol | 2014 | 10.1016/j.actao.2014.05.009 |
| *Tillandsia recurvata* | Plantae | Valverde; Bernal | Bol Soc Bot Mex | 2010 | 0366-2128 |
| *Tillandsia violacea* | Plantae | Mondrag√≥n; Ticktin | Conserv Biol | 2011 | 10.1111/j.1523-1739.2011.01691.x |
| *Vulpicida pinastri* | Fungi | Shriver; Cutler; Doak | Oecologia | 2012 | 10.1007/s00442-012-2301-4 |
| *Vriesea sanguinolenta* | Plantae | Zotz | Acta Oecol | 2005 | 10.1016/j.actao.2005.05.009 |
| *Asplenium adulterinum* | Plantae | Bucharov√°; M√ºnzbergov√°; T√°jek | Am J Bot | 2010 | 10.3732/ajb.0900351 |
| *Asplenium cuneifolium* | Plantae | Bucharov√°; M√ºnzbergov√°; T√°jek | Am J Bot | 2010 | 10.3732/ajb.0900351 |
| *Asplenium scolopendrium* | Plantae | Bremer; Jongejans | Popul Ecol | 2010 | 10.1007/s10144-009-0143-7 |
| *Actaea elata* | Plantae | Mayberry; Elle | Oecologia | 2010 | 10.1007/s00442-010-1809-8 |
| *Actaea spicata* | Plantae | Fr√∂borg; Eriksson | Can J Bot | 2003 | 10.1139/B03-099 |
| *Adenocarpus gibbsianus* | Plantae | Iriondo; Gim√©nez-Benavides; Albert; Lozano; Escudero | NA | 2009 | 978-84-8014-746-0 |
| *Agrimonia eupatoria* | Plantae | Kiviniemi | Plant Ecol | 2002 | 10.1111/j.1523-1739.2011.01691.x |
| *Alliaria petiolata* | Plantae | Evans; Davis; Raghu; Ragavendran; Landis; Schemske | Ecol Appl | 2012 | 10.1890/11-1291.1 |
| *Anarrhinum fruticosum* | Plantae | Iriondo; Gim√©nez-Benavides; Albert; Lozano; Escudero | NA | 2009 | 978-84-8014-746-0 |
| *Vitaliana primuliflora* | Plantae | Iriondo; Gim√©nez-Benavides; Albert; Lozano; Escudero | NA | 2009 | 978-84-8014-746-0 |
| *Anthericum ramosum* | Plantae | ƒåern√°; M√ºnzbergov√° | PLOS ONE | 2013 | 10.1371/journal.pone.0075563 |
| *Anthyllis vulneraria* | Plantae | Bastrenta; Lebreton; Thompson | J Ecol | 1995 | 10.2307/2261628 |
| *Anthyllis vulneraria* | Plantae | Marcante; Winkler; Erschbamer | Annals Bot | 2009 | 10.1093/aob/mcp047 |
| *Antirrhinum lopesianum* | Plantae | Iriondo; Gim√©nez-Benavides; Albert; Lozano; Escudero | NA | 2009 | 978-84-8014-746-0 |
| *Antirrhinum subbaeticum* | Plantae | Iriondo; Gim√©nez-Benavides; Albert; Lozano; Escudero | NA | 2009 | 978-84-8014-746-0 |
| *Boechera fecunda* | Plantae | Lesica; Shelly | Am J Bot | 1995 | 10.2307/2445615 |
| *Arenaria grandiflora subsp. bolosii* | Plantae | Iriondo; Gim√©nez-Benavides; Albert; Lozano; Escudero | NA | 2009 | 978-84-8014-746-0 |
| *Armeria merinoi* | Plantae | Iriondo; Gim√©nez-Benavides; Albert; Lozano; Escudero | NA | 2009 | 978-84-8014-746-0 |
| *Artemisia genipi* | Plantae | Marcante; Winkler; Erschbamer | Annals Bot | 2009 | 10.1093/aob/mcp047 |
| *Asarum canadense* | Plantae | Damman; Cain | J Ecol | 1998 | 10.1046/j.1365-2745.1998.00242.x |
| *Astragalus peckii* | Plantae | Martin; Meinke | Popul Ecol | 2012 | 10.1007/s10144-012-0318-5 |
| *Astragalus scaphoides* | Plantae | Lesica | Great Basin Nat | 1995 | NA |
| *Astragalus scaphoides* | Plantae | Crone; Lesica | Ecology | 2004 | 10.1890/03-0256 |
| *Astragalus tremolsianus* | Plantae | Iriondo; Gim√©nez-Benavides; Albert; Lozano; Escudero | NA | 2009 | 978-84-8014-746-0 |
| *Astragalus tyghensis* | Plantae | Kaye; Pyke | Ecology | 2003 | 10.1890/0012-9658(2003)084[1464:TEOSTO]2.0.CO;2 |
| *Boltonia decurrens* | Plantae | Smith; Caswell; Mettler-Cherry | Ecol Appl | 2005 | 10.1890/04-0434 |
| *Brassica insularis* | Plantae | Noel; Maurice; Mignot; Gl√©min; Carbonell; Justy; Guyot; Olivieri; Petit | Conserv Genet | 2010 | 10.1007/s10592-010-0056-1 |
| *Calathea ovandensis* | Plantae | Horvitz; Schemske | Ecol Monogr | 1995 | 10.2307/2937136 |
| *Calochortus lyallii* | Plantae | Miller; Antos; Allen | NA | 2004 | NA |
| *Calochortus lyallii* | Plantae | Miller; Antos; Allen | NA | 2004 | NA |
| *Calochortus macrocarpus* | Plantae | Miller; Antos; Allen | NA | 2004 | NA |
| *Carduus nutans* | Plantae | Jongejans; Sheppard; Shea | J Appl Ecol | 2006 | 10.1111/j.1365-2664.2006.01228.x |
| *Carex bigelowii* | Plantae | Carlsson; Callaghan | Oikos | 1991 | 10.2307/3544870 |
| *Carlina vulgaris* | Plantae | Lofgren; Eriksson; Lehtil√§ | Ann Bot Fenn | 2000 | NA |
| *Carlina vulgaris* | Plantae | Jongejans; Jorritsma-Wienk; Becker; Dost√°l; Mild√©n | J Ecol | 2010 | 10.1111/j.1365-2745.2009.01612.x |
| *Centaurea horrida* | Plantae | Pisanu; Farris; Filigheddu; Begona Garcia | Plant Ecol | 2012 | 10.1007/s11258-012-0110-9 |
| *Centaurea jacea* | Plantae | Jongejans; de Kroon | J Ecol | 2005 | 10.1111/j.1365-2745.2005.01003.x |
| *Chamaecrista lineata var. keyensis* | Plantae | Liu; Menges; Quintana-Ascencio | Ecol Appl | 2005 | 10.1890/03-5382 |
| *Chamaelirium luteum* | Plantae | Meagher; Antonovics | Ecology | 1982 | 10.2307/1940111 |
| *Cheirolophus metlesicsii* | Plantae | Iriondo; Gim√©nez-Benavides; Albert; Lozano; Escudero | NA | 2009 | 978-84-8014-746-0 |
| *Actaea elata* | Plantae | Kaye; Pyke | Ecology | 2003 | 10.1890/0012-9658(2003)084[1464:TEOSTO]2.0.CO;2 |
| *Actaea cordifolia* | Plantae | Cook | NA | 1993 | NA |
| *Cirsium dissectum* | Plantae | Jongejans; de Vere; de Kroon | Plant Ecol | 2008 | 10.1007/s11258-008-9397-y |
| *Cirsium pitcheri* | Plantae | Bell; Bowles; McEachern | NA | 2003 | 978-3-642-07869-9 |
| *Cirsium pitcheri* | Plantae | Ellis; Williams; Lesica; Bell; Bierzychudek; Bowles; Crone; Doak; Ehrl√©n; Ellis-Adam; McEachern; Ganesan; Latham; Luijten; Kaye; Knight; Menges; Morris; den Nijs; Oostermeijer; Quintana-Ascencio; Shelly; Stanley; Thorpe; Ticktin; Valverde; Weekley | Ecology | 2012 | 10.1890/11-1052.1 |
| *Cirsium pitcheri* | Plantae | Bell; Powell; Bowles | J Wildlife Manage | 2013 | 10.1002/jwmg.525 |
| *Cirsium pitcheri* | Plantae | Jolls; Marik; Hamz√©; Havens | Biol Conserv | 2015 | 10.1016/j.biocon.2015.04.006 |
| *Cirsium undulatum* | Plantae | Dalgleish; Koons; Adler | J Ecol | 2010 | 10.1111/j.1365-2745.2009.01585.x |
| *Cirsium vulgare* | Plantae | Bullock; Hill; Silvertown | J Ecol | 1994 | 10.2307/2261390 |
| *Cleistesiopsis bifaria* | Plantae | Wells; Willems | NA | 1991 | 90-5103-068-1 |
| *Cleistesiopsis divaricata* | Plantae | Wells; Willems | NA | 1991 | 90-5103-068-1 |
| *Colchicum autumnale* | Plantae | Winter; Jung; Eckstein; Otte; Donath; Kriechbaum | J Appl Ecol | 2014 | 10.1111/1365-2664.12217 |
| *Colchicum autumnale* | Plantae | Winter; Jung; Eckstein; Otte; Donath; Kriechbaum | J Appl Ecol | 2014 | 10.1111/1365-2664.12217 |
| *Corallorhiza trifida* | Plantae | Iriondo; Gim√©nez-Benavides; Albert; Lozano; Escudero | NA | 2009 | 978-84-8014-746-0 |
| *Cryptantha flava* | Plantae | Lucas; Forseth; Casper | J Ecol | 2008 | 10.1111/j.1365-2745.2007.01350.x |
| *Cypripedium calceolus* | Plantae | Garc√≠a; Go√±i; Guzman | Conserv Biol | 2010 | 10.1111/j.1523-1739.2010.01466.x |
| *Cypripedium calceolus* | Plantae | Garc√≠a; Go√±i; Guzman | Conserv Biol | 2010 | 10.1111/j.1523-1739.2010.01466.x |
| *Cypripedium fasciculatum* | Plantae | Thorpe; Stanley; Kayne; Latham | NA | 2011 | NA |
| *Danthonia sericea* | Plantae | Moloney | Ecology | 1988 | 10.2307/1941656 |
| *Dicentra canadensis* | Plantae | Lin; Miriti; Goodell | Ecol Evol | 2016 | 10.1002/ece3.2163 |
| *Dicerandra frutescens* | Plantae | Menges; Quintana-Ascencio; Weekley; Gaoue | Biol Conserv | 2006 | 10.1016/j.biocon.2005.08.002 |
| *Dipsacus fullonum* | Plantae | Werner; Caswell | Ecology | 1977 | 10.2307/1936930 |
| *Dorycnium spectabile* | Plantae | Iriondo; Gim√©nez-Benavides; Albert; Lozano; Escudero | NA | 2009 | 978-84-8014-746-0 |
| *Draba asterophora* | Plantae | Putnam | NA | 2013 | NA |
| *Dracocephalum austriacum* | Plantae | Andrello | NA | 2010 | NA |
| *Echinacea angustifolia* | Plantae | Dalgleish; Koons; Adler | J Ecol | 2010 | 10.1111/j.1365-2745.2009.01585.x |
| *Echinacea angustifolia* | Plantae | Hurlburt | NA | 1999 | NA |
| *Echinospartum ibericum subsp. algibicum* | Plantae | Iriondo; Gim√©nez-Benavides; Albert; Lozano; Escudero | NA | 2009 | 978-84-8014-746-0 |
| *Eriogonum longifolium var. gnaphalifolium* | Plantae | Satterthwaite; Menges; Quintana-Ascencio | Ecol Appl | 2002 | 10.1890/1051-0761(2002)012[1672:ASBPVI]2.0.CO;2 |
| *Erodium paularense* | Plantae | Iriondo; Gim√©nez-Benavides; Albert; Lozano; Escudero | NA | 2009 | 978-84-8014-746-0 |
| *Eryngium alpinum* | Plantae | Andrello; Bizoux; Barbet-Massin; Gaudeul; Nicol√®; Till-Bottraud | Biol Conserv | 2012 | 10.1016/j.biocon.2011.12.012 |
| *Eryngium cuneifolium* | Plantae | Menges; Quintana-Ascencio | Ecol Monogr | 2004 | 10.1890/03-4029 |
| *Eupatorium perfoliatum* | Plantae | Byers; Meagher | Ecol Appl | 1997 | 10.1890/1051-0761(1997)007[0519:ACODCI]2.0.CO;2 |
| *Eupatorium resinosum* | Plantae | Byers; Meagher | Ecol Appl | 1997 | 10.1890/1051-0761(1997)007[0519:ACODCI]2.0.CO;2 |
| *Oenothera coloradensis subsp. coloradensis* | Plantae | Floyd; Ranker | Int J Plant Sci | 1998 | 10.1086/297607 |
| *Gentiana pneumonanthe* | Plantae | Oostermeijer; Brugman; de Boer; den Nijs | J Ecol | 1996 | 10.2307/2261351 |
| *Geranium sylvaticum* | Plantae | Ramula; Toivonen; Mutikainen | Int J Plant Sci | 2007 | 10.1086/512040 |
| *Geum rivale* | Plantae | Kiviniemi | Plant Ecol | 2002 | 10.1111/j.1523-1739.2011.01691.x |
| *Pyrrocoma radiata* | Plantae | Kaye; Pyke | Ecology | 2003 | 10.1890/0012-9658(2003)084[1464:TEOSTO]2.0.CO;2 |
| *Stenaria nigricans* | Plantae | Dalgleish; Koons; Adler | J Ecol | 2010 | 10.1111/j.1365-2745.2009.01585.x |
| *Helianthemum polygonoides* | Plantae | Iriondo; Gim√©nez-Benavides; Albert; Lozano; Escudero | NA | 2009 | 978-84-8014-746-0 |
| *Helianthemum teneriffae* | Plantae | Iriondo; Gim√©nez-Benavides; Albert; Lozano; Escudero | NA | 2009 | 978-84-8014-746-0 |
| *Heliconia acuminata* | Plantae | Bruna | Ecology | 2003 | 10.1890/0012-9658(2003)084[0932:APPIFH]2.0.CO;2 |
| *Heliconia metallica* | Plantae | Schleuning; Huam√°n; Matthies | J Ecol | 2008 | 10.1111/j.1365-2745.2008.01416.x |
| *Hilaria mutica* | Plantae | Vega; Monta√±a | Plant Ecol | 2004 | 10.1023/B:VEGE.0000048094.21994.74 |
| *Horkelia congesta* | Plantae | Kaye; Benfield | NA | 2004 | NA |
| *Hydrastis canadensis* | Plantae | Sinclair | NA | 2002 | NA |
| *Tetraneuris herbacea* | Plantae | Campbell; Husband | Heredity | 2005 | 10.1038/sj.hdy.6800653 |
| *Hyparrhenia diplandra* | Plantae | Garnier; Dajoz | J Ecol | 2001 | 10.1890/0012-9658(2001)082[1720:ESOALV]2.0.CO;2 |
| *Hypericum cumulicola* | Plantae | Quintana-Ascencio; Menges; Weekley | Conserv Biol | 2003 | 10.1046/j.1523-1739.2003.01431.x |
| *Jurinea fontqueri* | Plantae | Iriondo; Gim√©nez-Benavides; Albert; Lozano; Escudero | NA | 2009 | 978-84-8014-746-0 |
| *Kosteletzkya pentacarpos* | Plantae | Pino; Pic√≥; Roa | Bot J Linn Soc | 2007 | 10.1111/j.1095-8339.2007.00628.x |
| *Kunkeliella subsucculenta* | Plantae | Iriondo; Gim√©nez-Benavides; Albert; Lozano; Escudero | NA | 2009 | 978-84-8014-746-0 |
| *Laserpitium longiradium* | Plantae | Iriondo; Gim√©nez-Benavides; Albert; Lozano; Escudero | NA | 2009 | 978-84-8014-746-0 |
| *Lathyrus Vernus* | Plantae | Ehrl√©n | J Ecol | 1995 | 10.2307/2261568 |
| *Leontopodium nivale subsp. alpinum* | Plantae | Keller; Vittoz | Alpine Bot | 2015 | 10.1007/s00035-014-0142-y |
| *Lepanthes rupestris* | Plantae | Tremblay; Ackerman | Biol J Linn Soc | 2001 | 10.1006/bijl.2000.0485 |
| *Lepanthes rupestris* | Plantae | Tremblay; McCarthy | PLOS ONE | 2014 | 10.1371/journal.pone.0102859 |
| *Lepidium davisii* | Plantae | Bernatus | NA | 1995 | NA |
| *Physaria ovalifolia* | Plantae | Dalgleish; Koons; Adler | J Ecol | 2010 | 10.1111/j.1365-2745.2009.01585.x |
| *Liatris scariosa* | Plantae | Ellis | Ecology | 2012 | 10.1890/11-1052.1 |
| *Limonium erectum* | Plantae | Iriondo; Gim√©nez-Benavides; Albert; Lozano; Escudero | NA | 2009 | 978-84-8014-746-0 |
| *Limonium geronense* | Plantae | Iriondo; Gim√©nez-Benavides; Albert; Lozano; Escudero | NA | 2009 | 978-84-8014-746-0 |
| *Limonium malacitanum* | Plantae | Iriondo; Gim√©nez-Benavides; Albert; Lozano; Escudero | NA | 2009 | 978-84-8014-746-0 |
| *Linum flavum* | Plantae | M√ºnzbergov√° | Plant Biology | 2013 | 10.1111/plb.12007 |
| *Linum tenuifolium* | Plantae | M√ºnzbergov√° | Plant Biology | 2013 | 10.1111/plb.12007 |
| *Lithospermum ruderale* | Plantae | Bricker; Maron | Ecology | 2012 | 10.1890/11-0948.1 |
| *Lomatium bradshawii* | Plantae | Kaye; Pyke | Ecology | 2003 | 10.1890/0012-9658(2003)084[1464:TEOSTO]2.0.CO;2 |
| *Lomatium bradshawii* | Plantae | Kaye; Pendergrass; Finley; Kauffman | Ecol Appl | 2001 | 10.1890/1051-0761(2001)011[1366:TEOFOT]2.0.CO;2 |
| *Lomatium cookii* | Plantae | Kaye; Pyke | Ecology | 2003 | 10.1890/0012-9658(2003)084[1464:TEOSTO]2.0.CO;2 |
| *Lotus arinagensis* | Plantae | Iriondo; Gim√©nez-Benavides; Albert; Lozano; Escudero | NA | 2009 | 978-84-8014-746-0 |
| *Lupinus lepidus* | Plantae | Bishop | NA | 1996 | NA |
| *Lupinus tidestromii* | Plantae | Dangremond; Knight | Ecology | 2010 | 10.1890/09-0418.1 |
| *Mimulus cardinalis* | Plantae | Angert | Ecology | 2006 | 10.1890/0012-9658(2006)87[2014:DOCAMP]2.0.CO;2 |
| *Molinia caerulea* | Plantae | Jacquemyn; Brys; Neubert | Ecol Appl | 2005 | 10.1890/04-1762 |
| *Narcissus pseudonarcissus* | Plantae | Barkham | J Ecol | 1980 | 10.2307/2259425 |
| *Neotinea ustulata* | Plantae | Shefferson; Tali | J Ecol | 2007 | 10.1111/j.1365-2745.2006.01195.x |
| *Oenothera deltoides* | Plantae | Thomson | Conserv Biol | 2005 | 10.1111/j.1523-1739.2005.004108.x |
| *Orchis purpurea* | Plantae | Jacquemyns; Brys; Jongejans | Ecology | 2010 | 10.1890/08-2321.1 |
| *Oxalis acetosella* | Plantae | Berg | Ecography | 2002 | 10.1034/j.1600-0587.2002.250211.x |
| *Oxytropis jabalambrensis* | Plantae | Iriondo; Gim√©nez-Benavides; Albert; Lozano; Escudero | NA | 2009 | 978-84-8014-746-0 |
| *Panax quinquefolius* | Plantae | Shahi | NA | 2007 | NA |
| *Panax quinquefolius* | Plantae | Charron; Gagnon | J Ecol | 1991 | 10.2307/2260724 |
| *Parolinia glabriuscula* | Plantae | Iriondo; Gim√©nez-Benavides; Albert; Lozano; Escudero | NA | 2009 | 978-84-8014-746-0 |
| *Paronychia jamesii* | Plantae | Dalgleish; Koons; Adler | J Ecol | 2010 | 10.1111/j.1365-2745.2009.01585.x |
| *Petrocoptis pyrenaica subsp. pseudoviscosa* | Plantae | Garc√≠a; Guzman; Go√±i | Biol Conserv | 2002 | 10.1016/S0006-3207(01)00113-6 |
| *Petrocoptis pyrenaica subsp. pseudoviscosa* | Plantae | Garc√≠a; Guzman; Go√±i | Biol Conserv | 2002 | 10.1016/S0006-3207(01)00113-6 |
| *Pimpinella saxifraga* | Plantae | Auestad; Rydgren; Jongejans; Kroon | Biol Conserv | 2010 | 10.1016/j.biocon.2009.12.037 |
| *Pinguicula ionantha* | Plantae | Kesler; Trusty; Hermann; Guyer | Oecologia | 2008 | 10.1007/s00442-008-1022-1 |
| *Plantago coronopus* | Plantae | Waite | J Ecol | 1984 | 10.2307/2259533 |
| *Plantago coronopus* | Plantae | Villellas; Ehrl√©n; Olesen; Braza; Garc√≠a | Ecography | 2013 | 10.1111/j.1600-0587.2012.07425.x |
| *Plantago coronopus* | Plantae | Villellas; Ehrl√©n; Olesen; Braza; Garc√≠a | Ecography | 2013 | 10.1111/j.1600-0587.2012.07425.x |
| *Plantago coronopus* | Plantae | Villellas; Ehrl√©n; Olesen; Braza; Garc√≠a | Ecography | 2013 | 10.1111/j.1600-0587.2012.07425.x |
| *Plantago coronopus* | Plantae | Villellas; Ehrl√©n; Olesen; Braza; Garc√≠a | Ecography | 2013 | 10.1111/j.1600-0587.2012.07425.x |
| *Plantago coronopus* | Plantae | Villellas; Ehrl√©n; Olesen; Braza; Garc√≠a | Ecography | 2013 | 10.1111/j.1600-0587.2012.07425.x |
| *Plantago media* | Plantae | Eriksson; Eriksson | J Veg Sci | 2000 | 10.2307/3236803 |
| *Poa alpina* | Plantae | Marcante; Winkler; Erschbamer | Annals Bot | 2009 | 10.1093/aob/mcp047 |
| *Polemonium van-bruntiae* | Plantae | Bermingham | Plant Ecol | 2010 | 10.1007/s11258-010-9762-5 |
| *Polygonella basiramia* | Plantae | Maliakal Witt | NA | 2004 | NA |
| *Potentilla anserina* | Plantae | Eriksson | J Ecol | 1988 | 10.2307/2260610 |
| *Primula elatior* | Plantae | Jacquemyn; Brys | Ecology | 2008 | 10.1890/07-1908.1 |
| *Primula farinosa* | Plantae | Lindborg; Ehrl√©n | Conserv Biol | 2002 | 10.1046/j.1523-1739.2002.00509.x |
| *Primula veris* | Plantae | Lehtil√§; Syrj√§nen; Leimu; Garc√≠a; Ehrl√©n | Conserv Biol | 2006 | 10.1111/j.1523-1739.2006.00368.x |
| *Primula veris* | Plantae | Lehtil√§; Syrj√§nen; Leimu; Garc√≠a; Ehrl√©n | Conserv Biol | 2006 | 10.1111/j.1523-1739.2006.00368.x |
| *Primula veris* | Plantae | Endels; Jacquemyn; Brys; Hermy | Plant Ecol | 2005 | 10.1007/s11258-004-0026-0 |
| *Primula vulgaris* | Plantae | Valverde; Silvertown | J Ecol | 1998 | 10.1046/j.1365-2745.1998.00280.x |
| *Primula vulgaris* | Plantae | Valdes; Garc√≠a; Garc√≠a; Ehrl√©n | Ecography | 2013 | 10.1111/j.1600-0587.2013.00216.x |
| *Pseudomisopates rivas-martinezii* | Plantae | Iriondo; Gim√©nez-Benavides; Albert; Lozano; Escudero | NA | 2009 | 978-84-8014-746-0 |
| *Psoralea tenuiflora* | Plantae | Dalgleish; Koons; Adler | J Ecol | 2010 | 10.1111/j.1365-2745.2009.01585.x |
| *Pyrrocoma radiata* | Plantae | Pfingsten | NA | 2013 | NA |
| *Ramonda myconi* | Plantae | Pic√≥; Riba | Plant Ecol | 2002 | 10.1023/A:1020310609348 |
| *Ranunculus peltatus* | Plantae | Idestam-Almquist | NA | 1998 | NA |
| *Ratibida columnifera* | Plantae | Dalgleish; Koons; Adler | J Ecol | 2010 | 10.1111/j.1365-2745.2009.01585.x |
| *Rubus praecox* | Plantae | Lambrecht-McDowell; Radosevich | Biol Invasions | 2005 | 10.1007/s10530-004-0870-9 |
| *Rumex rupestris* | Plantae | Iriondo; Gim√©nez-Benavides; Albert; Lozano; Escudero | NA | 2009 | 978-84-8014-746-0 |
| *Santolina melidensis* | Plantae | Iriondo; Gim√©nez-Benavides; Albert; Lozano; Escudero | NA | 2009 | 978-84-8014-746-0 |
| *Saponaria bellidifolia* | Plantae | Cserg≈ë; Moln√°r; Garc√≠a | Popul Ecol | 2011 | 10.1007/s10144-010-0249-y |
| *Sarcocapnos baetica* | Plantae | Salinas; Su√°rez; Blanca | Can J Bot | 2002 | 10.1139/b02-013 |
| *Sarcocapnos enneaphylla* | Plantae | Salinas; Su√°rez; Blanca | Can J Bot | 2002 | 10.1139/b02-013 |
| *Sarcocapnos pulcherrima* | Plantae | Salinas; Su√°rez; Blanca | Can J Bot | 2002 | 10.1139/b02-013 |
| *Sarracenia purpurea* | Plantae | Gotelli; Ellison | Ecol Appl | 2006 | 10.1890/04-0479 |
| *Saussurea medusa* | Plantae | Law; Salick; Knight | Plant Ecol | 2010 | 10.1007/s11258-010-9761-6 |
| *Saxifraga aizoides* | Plantae | Marcante; Winkler; Erschbamer | Annals Bot | 2009 | 10.1093/aob/mcp047 |
| *Saxifraga cotyledon* | Plantae | Dinnetz; Nilsson | Plant Ecol | 2002 | 10.1023/A:1015593311183 |
| *Silene douglasii var. oraria* | Plantae | Kephart; Paladino | Am J Bot | 1997 | 10.2307/2446079 |
| *Solidago mollis* | Plantae | Dalgleish; Koons; Adler | J Ecol | 2010 | 10.1111/j.1365-2745.2009.01585.x |
| *Sphaeralcea coccinea* | Plantae | Dalgleish; Koons; Adler | J Ecol | 2010 | 10.1111/j.1365-2745.2009.01585.x |
| *Sporobolus heterolepis* | Plantae | Dalgleish; Kula; Hartnett; Sandercock | Am J Bot | 2008 | 10.3732/ajb.2007277 |
| *Succisa pratensis* | Plantae | Wallin; Svensson | Folia Geobot | 2012 | 10.1007/s12224-012-9123-3 |
| *Succisa pratensis* | Plantae | Mild√©n | NA | 2005 | NA |
| *Taraxacum campylodes* | Plantae | Vavrek; McGraw; Yang | J Ecol | 1997 | 10.2307/2960501 |
| *Thelesperma megapotamicum* | Plantae | Dalgleish; Koons; Adler | J Ecol | 2010 | 10.1111/j.1365-2745.2009.01585.x |
| *Thymus vulgaris* | Plantae | Iriondo; Gim√©nez-Benavides; Albert; Lozano; Escudero | NA | 2009 | 978-84-8014-746-0 |
| *Trillium grandiflorum* | Plantae | Schmucki | NA | 2009 | NA |
| *Trillium ovatum* | Plantae | Ream | NA | 2011 | NA |
| *Trollius europaeus* | Plantae | Lemke; Salguero-G√≥mez | Popul Ecol | 2015 | 10.1007/s10144-015-0519-9 |
| *Trollius laxus* | Plantae | Scanga; Leopold | Biol Conserv | 2012 | 10.1016/j.biocon.2012.01.061 |
| *Trollius laxus* | Plantae | Scanga | Plant Ecol | 2014 | 10.1007/s11258-014-0344-9 |
| *Verbascum fontqueri* | Plantae | Iriondo; Gim√©nez-Benavides; Albert; Lozano; Escudero | NA | 2009 | 978-84-8014-746-0 |
| *Zea diploperennis* | Plantae | S√°nchez-Vel√°squez; Ezcurra; Martinez-Ramos; Alvarez-Buylla; Lorente | J Ecol | 2002 | 10.1046/j.1365-2745.2002.00702.x |
| *Alyxia stellata* | Plantae | Wong; Ticktin | Environ Conserv | 2014 | 10.1017/S0376892914000204 |
| *Machaerium cuspidatum* | Plantae | Nabe-Nielsen | J Trop Ecol | 2004 | 10.1017/S0266467404001609 |
| *Borassus aethiopum* | Plantae | Barot; Gignoux; Vuattoux | J Trop Ecol | 2000 | 10.1017/S0266467400001620 |
| *Ceratozamia mirandae* | Plantae | P√©rez-Farrera; Vovides; Octavio-Aguilar; Gonz√°lez-Astorga; Cruz-Rodr√≠guez; Hern√°ndez-Jonap√°; Villalobos-M√©ndez | Plant Ecol | 2006 | 10.1007/s11258-006-9135-2 |
| *Chamaedorea elegans* | Plantae | Valverde; Hern√°ndez-Apolinar; Mendoza-Amarom | J Sustain Forest | 2006 | 10.1300/J091v23n01_05 |
| *Chamaedorea radicalis* | Plantae | Endress; Gorchov; Robert; Noble | Ecol Appl | 2004 | 10.1890/02-5365 |
| *Chamaedorea radicalis* | Plantae | Berry; Gorchov; Endress; Stevens | Popul Ecol | 2008 | 10.1007/s10144-007-0067-z |
| *Dioon merolae* | Plantae | L√°zaro-Zerme√±o; Gonz√°lez-Espinosa; Mendoza; Martinez-Ramos; Quintana-Ascencio | Forest Ecol Manag | 2011 | 10.1016/j.foreco.2010.10.028 |
| *Eremospatha macrocarpa* | Plantae | Kouassi; Barot; Gignoux; Bi | J Trop Ecol | 2008 | 10.1017/S0266467408005312 |
| *Euterpe edulis* | Plantae | Silva-Matos; Freckleton; Watkinson | Ecology | 1999 | 10.1890/0012-9658(1999)080[2635:TRODDI]2.0.CO;2 |
| *Euterpe oleracea* | Plantae | Arango; Duque; Mu√±oz | Int J Trop Biol | 2010 | 10.15517/rbt.v58i1.5222 |
| *Geonoma pohliana subsp. weddelliana* | Plantae | Souza; Martins | Austral Ecol | 2006 | 10.1111/j.1442-9993.2006.01650.x |
| *Geonoma schottiana* | Plantae | Sampaio; Scariot | J Trop Ecol | 2010 | 10.1017/S0266467409990599 |
| *Iriartea deltoidea* | Plantae | Pinard | Biotropica | 1993 | 10.2307/2388974 |
| *Laccosperma secundiflorum* | Plantae | Kouassi; Barot; Gignoux; Bi | J Trop Ecol | 2008 | 10.1017/S0266467408005312 |
| *Pseudophoenix sargentii* | Plantae | Dur√°n; Franco | NA | 1992 | NA |
| *Sabal yapa* | Plantae | Pulido; Valverde; Caballero | J Trop Ecol | 2007 | 10.1017/S0266467406003877 |
| *Thrinax radiata* | Plantae | Olmsted; Alvarez-Buylla | Ecol Appl | 1995 | 10.2307/1942038 |
| *Zamia amblyphyllidia* | Plantae | Negron-Ortiz; Gorchov; Breckon | Int J Plant Sci | 1996 | 10.1086/297381 |
| *Acacia suaveolens* | Plantae | Warton; Wardle | Austral Ecol | 2003 | 10.1046/j.1442-9993.2003.01246.x |
| *Ardisia elliptica* | Plantae | Koop; Horvitz | Ecology | 2005 | 10.1890/04-1483 |
| *Atriplex acanthocarpa* | Plantae | Verhulst; Monta√±a; Mandujano; Franco | Oecologia | 2008 | 10.1007/s00442-008-0980-7 |
| *Atriplex canescens* | Plantae | Verhulst; Monta√±a; Mandujano; Franco | Oecologia | 2008 | 10.1007/s00442-008-0980-7 |
| *Clidemia hirta* | Plantae | DeWalt | Biol Invasions | 2006 | 10.1007/s10530-005-5277-8 |
| *Cytisus scoparius* | Plantae | Neubert; Parker | Risk Anal | 2004 | 10.1111/j.0272-4332.2004.00481.x |
| *Fumana procumbens* | Plantae | Bengtsson | J Ecol | 1993 | 10.2307/2261672 |
| *Gardenia actinocarpa* | Plantae | Osunkoya | Biol Conserv | 2003 | 10.1016/S0006-3207(02)00417-2 |
| *Helianthemum juliae* | Plantae | Marrero-G√≥mez; Oostermeijer; Carqu√©-√Ålamo; Ba√±ares-Baudet | Biol Conserv | 2007 | 10.1016/j.biocon.2007.01.010 |
| *Persoonia bargoensis* | Plantae | McKenna | NA | 2007 | NA |
| *Persoonia glaucescens* | Plantae | McKenna | NA | 2007 | NA |
| *Purshia subintegra* | Plantae | Maschinski; Baggs; Quintana-Ascencio; Menges | Conserv Biol | 2006 | 10.1111/j.1523-1739.2006.00272.x |
| *Rosmarinus tomentosus* | Plantae | Iriondo; Gim√©nez-Benavides; Albert; Lozano; Escudero | NA | 2009 | 978-84-8014-746-0 |
| *Tetramolopium arenarium* | Plantae | Aplet; Laven; Shaw | Nat Area J | 1994 | NA |
| *Vella pseudocytisus subsp. paui* | Plantae | Iriondo; Gim√©nez-Benavides; Albert; Lozano; Escudero | NA | 2009 | 978-84-8014-746-0 |
| *Vella pseudocytisus subsp. paui* | Plantae | Iriondo; Gim√©nez-Benavides; Albert; Lozano; Escudero | NA | 2009 | 978-84-8014-746-0 |
| *Astrophytum asterias* | Plantae | Martinez-Avalos | NA | 2007 | NA |
| *Astrophytum ornatum* | Plantae | Zepeda-Martinez; Mandujano; Mandujano; Golubov | J Arid Environ | 2013 | 10.1016/j.jaridenv.2012.08.006 |
| *Escobaria robbinsorum* | Plantae | Schmalzel; Reichenbacher; Rutman | Madrono | 1995 | NA |
| *Escontria chiotilla* | Plantae | Ortega-Baes | NA | 2001 | NA |
| *Euphorbia fontqueriana* | Plantae | Iriondo; Gim√©nez-Benavides; Albert; Lozano; Escudero | NA | 2009 | 978-84-8014-746-0 |
| *Mammillaria crucigera* | Plantae | Contreras; Valverde | J Arid Environ | 2002 | 10.1006/jare.2001.0926 |
| *Mammillaria gaumeri* | Plantae | Ferrer; Dur√°n; M√©ndez; Dorantes; Dzib | Bol Soc Bot Mex | 2011 | NA |
| *Mammillaria hernandezii* | Plantae | Rodriguez Ortega | NA | 2008 | NA |
| *Mammillaria huitzilopochtli* | Plantae | Flores Mart√≠nez; Manzanero-Medino; Golubov; Monta√±a; Mandujano | Plant Ecol | 2010 | 10.1007/s11258-010-9737-6 |
| *Mammillaria huitzilopochtli* | Plantae | Flores Mart√≠nez | NA | 2010 | NA |
| *Mammillaria magnimamma* | Plantae | Valverde; Quijas; Lopez-Villavicencio; Castillo | Plant Ecol | 2004 | 10.1023/B:VEGE.0000021662.78634.de |
| *Mammillaria napina* | Plantae | Rodriguez Ortega | NA | 2008 | NA |
| *Mammillaria solisioides* | Plantae | Rodriguez Ortega | NA | 2008 | NA |
| *Neobuxbaumia macrocephala* | Plantae | Esparza-Olgu√≠n; Valverde; Mandujano | Popul Ecol | 2005 | 10.1007/s10144-005-0230-3 |
| *Neobuxbaumia macrocephala* | Plantae | Esparza Olgu√≠n | NA | 2005 | NA |
| *Neobuxbaumia mezcalaensis* | Plantae | Esparza-Olgu√≠n; Valverde; Mandujano | Popul Ecol | 2005 | 10.1007/s10144-005-0230-3 |
| *Neobuxbaumia mezcalaensis* | Plantae | Esparza Olgu√≠n | NA | 2005 | NA |
| *Neobuxbaumia polylopha* | Plantae | Arroyo-Cosultchi; Golubov; Mandujano | Acta Oecol | 2016 | 10.1016/j.actao.2016.01.006 |
| *Neobuxbaumia tetetzo* | Plantae | Esparza-Olgu√≠n; Valverde; Mandujano | Popul Ecol | 2005 | 10.1007/s10144-005-0230-3 |
| *Neobuxbaumia tetetzo* | Plantae | Esparza Olgu√≠n | NA | 2005 | NA |
| *Opuntia macrorhiza* | Plantae | Haridas; Keeler; Tenhumberg | Ecology | 2015 | 10.1890/13-1984.1 |
| *Opuntia rastrera* | Plantae | Mandujano; Monta√±a; Franco; Golubov; Flores-Mart√≠nez | Ecology | 2001 | 10.2307/2679864 |
| *Pediocactus bradyi* | Plantae | Shryock; Esque; Hughes | Am J Bot | 2014 | 10.3732/ajb.1400035 |
| *Pterocereus gaumeri* | Plantae | M√©ndez; Dur√°n; Olmsted | Biotropica | 2004 | 10.1646/1601 |
| *Stenocereus eruca* | Plantae | Clark-Tapia | NA | 2004 | NA |
| *Acer saccharum* | Plantae | Lin; Augspurger | Forest Ecol Manag | 2008 | 10.1016/j.foreco.2008.02.040 |
| *Aesculus turbinata* | Plantae | Kaneko; Takada; Kawano | Plant Spec Biol | 1999 | 10.1046/j.1442-1984.1999.00007.x |
| *Alnus incana subsp. rugosa* | Plantae | Huenneke; Marks | Ecology | 1987 | 10.2307/1939207 |
| *Bursera glabrifolia* | Plantae | Hern√°ndez-Apolinar; Valverde; Purata | Forest Ecol Manag | 2006 | 10.1016/j.foreco.2005.10.072 |
| *Castanea dentata* | Plantae | Davelos; Jarosz | J Ecol | 2004 | 10.1111/j.0022-0477.2004.00907.x |
| *Fagus grandifolia* | Plantae | da Silva Batista; Platt; Macchiavelli | Ecology | 1998 | 10.2307/176863 |
| *Khaya senegalensis* | Plantae | Gaoue; Ticktin | Conserv Biol | 2010 | 10.1111/j.1523-1739.2009.01345.x |
| *Magnolia macrophylla var. dealbata* | Plantae | S√°nchez-Vel√°squez; Pineda-L√≥pez | Popul Ecol | 2010 | 10.1007/s10144-009-0161-5 |
| *Manilkara zapota* | Plantae | Cruz-Rodr√≠guez; Lopez-Villavicencio; Valverde | J Trop Ecol | 2009 | 10.1017/S0266467408005713 |
| *Phyllanthus emblica* | Plantae | Ticktin; Ganesan; Paramesha; Setty | J Appl Ecol | 2012 | 10.1111/j.1365-2664.2012.02156.x |
| *Phyllanthus emblica* | Plantae | Ellis; Williams; Lesica; Bell; Bierzychudek; Bowles; Crone; Doak; Ehrl√©n; Ellis-Adam; McEachern; Ganesan; Latham; Luijten; Kaye; Knight; Menges; Morris; den Nijs; Oostermeijer; Quintana-Ascencio; Shelly; Stanley; Thorpe; Ticktin; Valverde; Weekley | Ecology | 2012 | 10.1890/11-1052.1 |
| *Phyllanthus indofischeri* | Plantae | Ticktin; Ganesan; Paramesha; Setty | J Appl Ecol | 2012 | 10.1111/j.1365-2664.2012.02156.x |
| *Pinus lambertiana* | Plantae | van Mantgem; Stephenson | J Ecol | 2005 | 10.1111/j.1365-2745.2005.01007.x |
| *Pinus lambertiana* | Plantae | Maloney; Vogler; Eckert; Jensen; Neale | Forest Ecol Manag | 2011 | 10.1016/j.foreco.2011.05.011 |
| *Pinus nigra* | Plantae | Buckley; Brockerhoff; Langer; Ledgard; North; Rees | J Appl Ecol | 2005 | 10.1111/j.1365-2664.2005.01100.x |
| *Pinus strobus* | Plantae | M√ºnzbergov√°; Hadincov√°; Wild; Kindlmannov√° | PLOS ONE | 2013 | 10.1371/journal.pone.0056953 |
| *Prioria copaifera* | Plantae | Condit | Forest Ecol Manag | 1993 | 10.1016/0378-1127(93)90045-O |
| *Prosopis laevigata* | Plantae | Bernal | NA | 2004 | NA |
| *Prunus africana* | Plantae | Stewart | NA | 2001 | NA |
| *Rhododendron ponticum* | Plantae | Salguero-G√≥mez | NA | 2004 | NA |
| *Rhododendron ponticum* | Plantae | Salguero-G√≥mez | NA | 2004 | NA |
| *Sapium sebiferum* | Plantae | Renne | NA | 2001 | NA |
| *Shorea leprosula* | Plantae | Chen; Visser; Jongejans; van Breugel; Zuidema; Kassim; de Kroon | J Ecol | 2011 | 10.1111/j.1365-2745.2011.01825.x |
| *Styrax obassis* | Plantae | Abe; Nokashizuka; Tanoka | J Veg Sci | 1998 | 10.2307/3237044 |
| *Taxus floridana* | Plantae | Kwit; Horvitz; Platt | Conserv Biol | 2004 | 10.1111/j.1523-1739.2004.00567.x |
| *Tsuga canadensis* | Plantae | Lamar; McGraw | Forest Ecol Manag | 2005 | 10.1016/j.foreco.2005.02.056 |
| *Ziziphus jujuba* | Plantae | Zull; Lawes; Cacho | Environ Modell Softw | 2015 | 10.1016/j.envsoft.2015.10.026 |
| *Oeceoclades maculata* | Plantae | River√≥n-Gir√≥; Ravent√≥s; Damon; Garc√≠a-Gonz√°lez; M√∫jica | Biol Invasions | 2019 | 10.1007/s10530-019-01945-7 |
| *Zea diploperennis* | Plantae | S√°nchez-Vel√°squez; Ezcurra; Martinez-Ramos; Alvarez-Buylla; Lorente | J Ecol | 2002 | 10.1046/j.1365-2745.2002.00702.x |
| *Alyxia stellata* | Plantae | Wong; Ticktin | Environ Conserv | 2014 | 10.1017/S0376892914000204 |
| *Machaerium cuspidatum* | Plantae | Nabe-Nielsen | J Trop Ecol | 2004 | 10.1017/S0266467404001609 |
| *Borassus aethiopum* | Plantae | Barot; Gignoux; Vuattoux | J Trop Ecol | 2000 | 10.1017/S0266467400001620 |
| *Ceratozamia mirandae* | Plantae | P√©rez-Farrera; Vovides; Octavio-Aguilar; Gonz√°lez-Astorga; Cruz-Rodr√≠guez; Hern√°ndez-Jonap√°; Villalobos-M√©ndez | Plant Ecol | 2006 | 10.1007/s11258-006-9135-2 |
| *Chamaedorea elegans* | Plantae | Valverde; Hern√°ndez-Apolinar; Mendoza-Amarom | J Sustain Forest | 2006 | 10.1300/J091v23n01_05 |
| *Chamaedorea radicalis* | Plantae | Endress; Gorchov; Robert; Noble | Ecol Appl | 2004 | 10.1890/02-5365 |
| *Chamaedorea radicalis* | Plantae | Berry; Gorchov; Endress; Stevens | Popul Ecol | 2008 | 10.1007/s10144-007-0067-z |
| *Dioon merolae* | Plantae | L√°zaro-Zerme√±o; Gonz√°lez-Espinosa; Mendoza; Martinez-Ramos; Quintana-Ascencio | Forest Ecol Manag | 2011 | 10.1016/j.foreco.2010.10.028 |
| *Eremospatha macrocarpa* | Plantae | Kouassi; Barot; Gignoux; Bi | J Trop Ecol | 2008 | 10.1017/S0266467408005312 |
| *Euterpe edulis* | Plantae | Silva-Matos; Freckleton; Watkinson | Ecology | 1999 | 10.1890/0012-9658(1999)080[2635:TRODDI]2.0.CO;2 |
| *Euterpe oleracea* | Plantae | Arango; Duque; Mu√±oz | Int J Trop Biol | 2010 | 10.15517/rbt.v58i1.5222 |
| *Geonoma pohliana subsp. weddelliana* | Plantae | Souza; Martins | Austral Ecol | 2006 | 10.1111/j.1442-9993.2006.01650.x |
| *Geonoma schottiana* | Plantae | Sampaio; Scariot | J Trop Ecol | 2010 | 10.1017/S0266467409990599 |
| *Iriartea deltoidea* | Plantae | Pinard | Biotropica | 1993 | 10.2307/2388974 |
| *Laccosperma secundiflorum* | Plantae | Kouassi; Barot; Gignoux; Bi | J Trop Ecol | 2008 | 10.1017/S0266467408005312 |
| *Pseudophoenix sargentii* | Plantae | Dur√°n; Franco | NA | 1992 | NA |
| *Sabal yapa* | Plantae | Pulido; Valverde; Caballero | J Trop Ecol | 2007 | 10.1017/S0266467406003877 |
| *Thrinax radiata* | Plantae | Olmsted; Alvarez-Buylla | Ecol Appl | 1995 | 10.2307/1942038 |
| *Zamia amblyphyllidia* | Plantae | Negron-Ortiz; Gorchov; Breckon | Int J Plant Sci | 1996 | 10.1086/297381 |
| *Acacia suaveolens* | Plantae | Warton; Wardle | Austral Ecol | 2003 | 10.1046/j.1442-9993.2003.01246.x |
| *Ardisia elliptica* | Plantae | Koop; Horvitz | Ecology | 2005 | 10.1890/04-1483 |
| *Atriplex acanthocarpa* | Plantae | Verhulst; Monta√±a; Mandujano; Franco | Oecologia | 2008 | 10.1007/s00442-008-0980-7 |
| *Atriplex canescens* | Plantae | Verhulst; Monta√±a; Mandujano; Franco | Oecologia | 2008 | 10.1007/s00442-008-0980-7 |
| *Clidemia hirta* | Plantae | DeWalt | Biol Invasions | 2006 | 10.1007/s10530-005-5277-8 |
| *Cytisus scoparius* | Plantae | Neubert; Parker | Risk Anal | 2004 | 10.1111/j.0272-4332.2004.00481.x |
| *Fumana procumbens* | Plantae | Bengtsson | J Ecol | 1993 | 10.2307/2261672 |
| *Gardenia actinocarpa* | Plantae | Osunkoya | Biol Conserv | 2003 | 10.1016/S0006-3207(02)00417-2 |
| *Helianthemum juliae* | Plantae | Marrero-G√≥mez; Oostermeijer; Carqu√©-√Ålamo; Ba√±ares-Baudet | Biol Conserv | 2007 | 10.1016/j.biocon.2007.01.010 |
| *Persoonia bargoensis* | Plantae | McKenna | NA | 2007 | NA |
| *Persoonia glaucescens* | Plantae | McKenna | NA | 2007 | NA |
| *Purshia subintegra* | Plantae | Maschinski; Baggs; Quintana-Ascencio; Menges | Conserv Biol | 2006 | 10.1111/j.1523-1739.2006.00272.x |
| *Rosmarinus tomentosus* | Plantae | Iriondo; Gim√©nez-Benavides; Albert; Lozano; Escudero | NA | 2009 | 978-84-8014-746-0 |
| *Tetramolopium arenarium* | Plantae | Aplet; Laven; Shaw | Nat Area J | 1994 | NA |
| *Vella pseudocytisus subsp. paui* | Plantae | Iriondo; Gim√©nez-Benavides; Albert; Lozano; Escudero | NA | 2009 | 978-84-8014-746-0 |
| *Vella pseudocytisus subsp. paui* | Plantae | Iriondo; Gim√©nez-Benavides; Albert; Lozano; Escudero | NA | 2009 | 978-84-8014-746-0 |
| *Astrophytum asterias* | Plantae | Martinez-Avalos | NA | 2007 | NA |
| *Astrophytum ornatum* | Plantae | Zepeda-Martinez; Mandujano; Mandujano; Golubov | J Arid Environ | 2013 | 10.1016/j.jaridenv.2012.08.006 |
| *Escobaria robbinsorum* | Plantae | Schmalzel; Reichenbacher; Rutman | Madrono | 1995 | NA |
| *Escontria chiotilla* | Plantae | Ortega-Baes | NA | 2001 | NA |
| *Euphorbia fontqueriana* | Plantae | Iriondo; Gim√©nez-Benavides; Albert; Lozano; Escudero | NA | 2009 | 978-84-8014-746-0 |
| *Mammillaria crucigera* | Plantae | Contreras; Valverde | J Arid Environ | 2002 | 10.1006/jare.2001.0926 |
| *Mammillaria gaumeri* | Plantae | Ferrer; Dur√°n; M√©ndez; Dorantes; Dzib | Bol Soc Bot Mex | 2011 | NA |
| *Mammillaria hernandezii* | Plantae | Rodriguez Ortega | NA | 2008 | NA |
| *Mammillaria huitzilopochtli* | Plantae | Flores Mart√≠nez; Manzanero-Medino; Golubov; Monta√±a; Mandujano | Plant Ecol | 2010 | 10.1007/s11258-010-9737-6 |
| *Mammillaria huitzilopochtli* | Plantae | Flores Mart√≠nez | NA | 2010 | NA |
| *Mammillaria magnimamma* | Plantae | Valverde; Quijas; Lopez-Villavicencio; Castillo | Plant Ecol | 2004 | 10.1023/B:VEGE.0000021662.78634.de |
| *Mammillaria napina* | Plantae | Rodriguez Ortega | NA | 2008 | NA |
| *Mammillaria solisioides* | Plantae | Rodriguez Ortega | NA | 2008 | NA |
| *Neobuxbaumia macrocephala* | Plantae | Esparza-Olgu√≠n; Valverde; Mandujano | Popul Ecol | 2005 | 10.1007/s10144-005-0230-3 |
| *Neobuxbaumia macrocephala* | Plantae | Esparza Olgu√≠n | NA | 2005 | NA |
| *Neobuxbaumia mezcalaensis* | Plantae | Esparza-Olgu√≠n; Valverde; Mandujano | Popul Ecol | 2005 | 10.1007/s10144-005-0230-3 |
| *Neobuxbaumia mezcalaensis* | Plantae | Esparza Olgu√≠n | NA | 2005 | NA |
| *Neobuxbaumia polylopha* | Plantae | Arroyo-Cosultchi; Golubov; Mandujano | Acta Oecol | 2016 | 10.1016/j.actao.2016.01.006 |
| *Neobuxbaumia tetetzo* | Plantae | Esparza-Olgu√≠n; Valverde; Mandujano | Popul Ecol | 2005 | 10.1007/s10144-005-0230-3 |
| *Neobuxbaumia tetetzo* | Plantae | Esparza Olgu√≠n | NA | 2005 | NA |
| *Opuntia macrorhiza* | Plantae | Haridas; Keeler; Tenhumberg | Ecology | 2015 | 10.1890/13-1984.1 |
| *Opuntia rastrera* | Plantae | Mandujano; Monta√±a; Franco; Golubov; Flores-Mart√≠nez | Ecology | 2001 | 10.2307/2679864 |
| *Pediocactus bradyi* | Plantae | Shryock; Esque; Hughes | Am J Bot | 2014 | 10.3732/ajb.1400035 |
| *Pterocereus gaumeri* | Plantae | M√©ndez; Dur√°n; Olmsted | Biotropica | 2004 | 10.1646/1601 |
| *Stenocereus eruca* | Plantae | Clark-Tapia | NA | 2004 | NA |
| *Acer saccharum* | Plantae | Lin; Augspurger | Forest Ecol Manag | 2008 | 10.1016/j.foreco.2008.02.040 |
| *Aesculus turbinata* | Plantae | Kaneko; Takada; Kawano | Plant Spec Biol | 1999 | 10.1046/j.1442-1984.1999.00007.x |
| *Alnus incana subsp. rugosa* | Plantae | Huenneke; Marks | Ecology | 1987 | 10.2307/1939207 |
| *Bursera glabrifolia* | Plantae | Hern√°ndez-Apolinar; Valverde; Purata | Forest Ecol Manag | 2006 | 10.1016/j.foreco.2005.10.072 |
| *Castanea dentata* | Plantae | Davelos; Jarosz | J Ecol | 2004 | 10.1111/j.0022-0477.2004.00907.x |
| *Fagus grandifolia* | Plantae | da Silva Batista; Platt; Macchiavelli | Ecology | 1998 | 10.2307/176863 |
| *Khaya senegalensis* | Plantae | Gaoue; Ticktin | Conserv Biol | 2010 | 10.1111/j.1523-1739.2009.01345.x |
| *Magnolia macrophylla var. dealbata* | Plantae | S√°nchez-Vel√°squez; Pineda-L√≥pez | Popul Ecol | 2010 | 10.1007/s10144-009-0161-5 |
| *Manilkara zapota* | Plantae | Cruz-Rodr√≠guez; Lopez-Villavicencio; Valverde | J Trop Ecol | 2009 | 10.1017/S0266467408005713 |
| *Phyllanthus emblica* | Plantae | Ticktin; Ganesan; Paramesha; Setty | J Appl Ecol | 2012 | 10.1111/j.1365-2664.2012.02156.x |
| *Phyllanthus emblica* | Plantae | Ellis; Williams; Lesica; Bell; Bierzychudek; Bowles; Crone; Doak; Ehrl√©n; Ellis-Adam; McEachern; Ganesan; Latham; Luijten; Kaye; Knight; Menges; Morris; den Nijs; Oostermeijer; Quintana-Ascencio; Shelly; Stanley; Thorpe; Ticktin; Valverde; Weekley | Ecology | 2012 | 10.1890/11-1052.1 |
| *Phyllanthus indofischeri* | Plantae | Ticktin; Ganesan; Paramesha; Setty | J Appl Ecol | 2012 | 10.1111/j.1365-2664.2012.02156.x |
| *Pinus lambertiana* | Plantae | van Mantgem; Stephenson | J Ecol | 2005 | 10.1111/j.1365-2745.2005.01007.x |
| *Pinus lambertiana* | Plantae | Maloney; Vogler; Eckert; Jensen; Neale | Forest Ecol Manag | 2011 | 10.1016/j.foreco.2011.05.011 |
| *Pinus nigra* | Plantae | Buckley; Brockerhoff; Langer; Ledgard; North; Rees | J Appl Ecol | 2005 | 10.1111/j.1365-2664.2005.01100.x |
| *Pinus strobus* | Plantae | M√ºnzbergov√°; Hadincov√°; Wild; Kindlmannov√° | PLOS ONE | 2013 | 10.1371/journal.pone.0056953 |
| *Prioria copaifera* | Plantae | Condit | Forest Ecol Manag | 1993 | 10.1016/0378-1127(93)90045-O |
| *Prosopis laevigata* | Plantae | Bernal | NA | 2004 | NA |
| *Prunus africana* | Plantae | Stewart | NA | 2001 | NA |
| *Rhododendron ponticum* | Plantae | Salguero-G√≥mez | NA | 2004 | NA |
| *Rhododendron ponticum* | Plantae | Salguero-G√≥mez | NA | 2004 | NA |
| *Sapium sebiferum* | Plantae | Renne | NA | 2001 | NA |
| *Shorea leprosula* | Plantae | Chen; Visser; Jongejans; van Breugel; Zuidema; Kassim; de Kroon | J Ecol | 2011 | 10.1111/j.1365-2745.2011.01825.x |
| *Styrax obassis* | Plantae | Abe; Nokashizuka; Tanoka | J Veg Sci | 1998 | 10.2307/3237044 |
| *Taxus floridana* | Plantae | Kwit; Horvitz; Platt | Conserv Biol | 2004 | 10.1111/j.1523-1739.2004.00567.x |
| *Tsuga canadensis* | Plantae | Lamar; McGraw | Forest Ecol Manag | 2005 | 10.1016/j.foreco.2005.02.056 |
| *Ziziphus jujuba* | Plantae | Zull; Lawes; Cacho | Environ Modell Softw | 2015 | 10.1016/j.envsoft.2015.10.026 |
| *Oeceoclades maculata* | Plantae | River√≥n-Gir√≥; Ravent√≥s; Damon; Garc√≠a-Gonz√°lez; M√∫jica | Biol Invasions | 2019 | 10.1007/s10530-019-01945-7 |
| *Neobuxbaumia macrocephala* | Plantae | Esparza-Olgu√≠n; Valverde; Mandujano | Popul Ecol | 2005 | 10.1007/s10144-005-0230-3 |
| *Neobuxbaumia macrocephala* | Plantae | Esparza Olgu√≠n | NA | 2005 | NA |
| *Neobuxbaumia mezcalaensis* | Plantae | Esparza-Olgu√≠n; Valverde; Mandujano | Popul Ecol | 2005 | 10.1007/s10144-005-0230-3 |
| *Neobuxbaumia mezcalaensis* | Plantae | Esparza Olgu√≠n | NA | 2005 | NA |
| *Neobuxbaumia polylopha* | Plantae | Arroyo-Cosultchi; Golubov; Mandujano | Acta Oecol | 2016 | 10.1016/j.actao.2016.01.006 |
| *Neobuxbaumia tetetzo* | Plantae | Esparza-Olgu√≠n; Valverde; Mandujano | Popul Ecol | 2005 | 10.1007/s10144-005-0230-3 |
| *Neobuxbaumia tetetzo* | Plantae | Esparza Olgu√≠n | NA | 2005 | NA |
| *Opuntia macrorhiza* | Plantae | Haridas; Keeler; Tenhumberg | Ecology | 2015 | 10.1890/13-1984.1 |
| *Opuntia rastrera* | Plantae | Mandujano; Monta√±a; Franco; Golubov; Flores-Mart√≠nez | Ecology | 2001 | 10.2307/2679864 |
| *Pediocactus bradyi* | Plantae | Shryock; Esque; Hughes | Am J Bot | 2014 | 10.3732/ajb.1400035 |
| *Pterocereus gaumeri* | Plantae | M√©ndez; Dur√°n; Olmsted | Biotropica | 2004 | 10.1646/1601 |
| *Stenocereus eruca* | Plantae | Clark-Tapia | NA | 2004 | NA |
| *Acer saccharum* | Plantae | Lin; Augspurger | Forest Ecol Manag | 2008 | 10.1016/j.foreco.2008.02.040 |
| *Aesculus turbinata* | Plantae | Kaneko; Takada; Kawano | Plant Spec Biol | 1999 | 10.1046/j.1442-1984.1999.00007.x |
| *Alnus incana subsp. rugosa* | Plantae | Huenneke; Marks | Ecology | 1987 | 10.2307/1939207 |
| *Bursera glabrifolia* | Plantae | Hern√°ndez-Apolinar; Valverde; Purata | Forest Ecol Manag | 2006 | 10.1016/j.foreco.2005.10.072 |
| *Castanea dentata* | Plantae | Davelos; Jarosz | J Ecol | 2004 | 10.1111/j.0022-0477.2004.00907.x |
| *Fagus grandifolia* | Plantae | da Silva Batista; Platt; Macchiavelli | Ecology | 1998 | 10.2307/176863 |
| *Khaya senegalensis* | Plantae | Gaoue; Ticktin | Conserv Biol | 2010 | 10.1111/j.1523-1739.2009.01345.x |
| *Magnolia macrophylla var. dealbata* | Plantae | S√°nchez-Vel√°squez; Pineda-L√≥pez | Popul Ecol | 2010 | 10.1007/s10144-009-0161-5 |
| *Manilkara zapota* | Plantae | Cruz-Rodr√≠guez; Lopez-Villavicencio; Valverde | J Trop Ecol | 2009 | 10.1017/S0266467408005713 |
| *Phyllanthus emblica* | Plantae | Ticktin; Ganesan; Paramesha; Setty | J Appl Ecol | 2012 | 10.1111/j.1365-2664.2012.02156.x |
| *Phyllanthus emblica* | Plantae | Ellis; Williams; Lesica; Bell; Bierzychudek; Bowles; Crone; Doak; Ehrl√©n; Ellis-Adam; McEachern; Ganesan; Latham; Luijten; Kaye; Knight; Menges; Morris; den Nijs; Oostermeijer; Quintana-Ascencio; Shelly; Stanley; Thorpe; Ticktin; Valverde; Weekley | Ecology | 2012 | 10.1890/11-1052.1 |
| *Phyllanthus indofischeri* | Plantae | Ticktin; Ganesan; Paramesha; Setty | J Appl Ecol | 2012 | 10.1111/j.1365-2664.2012.02156.x |
| *Pinus lambertiana* | Plantae | van Mantgem; Stephenson | J Ecol | 2005 | 10.1111/j.1365-2745.2005.01007.x |
| *Pinus lambertiana* | Plantae | Maloney; Vogler; Eckert; Jensen; Neale | Forest Ecol Manag | 2011 | 10.1016/j.foreco.2011.05.011 |
| *Pinus nigra* | Plantae | Buckley; Brockerhoff; Langer; Ledgard; North; Rees | J Appl Ecol | 2005 | 10.1111/j.1365-2664.2005.01100.x |
| *Pinus strobus* | Plantae | M√ºnzbergov√°; Hadincov√°; Wild; Kindlmannov√° | PLOS ONE | 2013 | 10.1371/journal.pone.0056953 |
| *Prioria copaifera* | Plantae | Condit | Forest Ecol Manag | 1993 | 10.1016/0378-1127(93)90045-O |
| *Prosopis laevigata* | Plantae | Bernal | NA | 2004 | NA |
| *Prunus africana* | Plantae | Stewart | NA | 2001 | NA |
| *Rhododendron ponticum* | Plantae | Salguero-G√≥mez | NA | 2004 | NA |
| *Rhododendron ponticum* | Plantae | Salguero-G√≥mez | NA | 2004 | NA |
| *Sapium sebiferum* | Plantae | Renne | NA | 2001 | NA |
| *Shorea leprosula* | Plantae | Chen; Visser; Jongejans; van Breugel; Zuidema; Kassim; de Kroon | J Ecol | 2011 | 10.1111/j.1365-2745.2011.01825.x |
| *Styrax obassis* | Plantae | Abe; Nokashizuka; Tanoka | J Veg Sci | 1998 | 10.2307/3237044 |
| *Taxus floridana* | Plantae | Kwit; Horvitz; Platt | Conserv Biol | 2004 | 10.1111/j.1523-1739.2004.00567.x |
| *Tsuga canadensis* | Plantae | Lamar; McGraw | Forest Ecol Manag | 2005 | 10.1016/j.foreco.2005.02.056 |
| *Ziziphus jujuba* | Plantae | Zull; Lawes; Cacho | Environ Modell Softw | 2015 | 10.1016/j.envsoft.2015.10.026 |
| *Oeceoclades maculata* | Plantae | River√≥n-Gir√≥; Ravent√≥s; Damon; Garc√≠a-Gonz√°lez; M√∫jica | Biol Invasions | 2019 | 10.1007/s10530-019-01945-7 |
| *Fagus grandifolia* | Plantae | da Silva Batista; Platt; Macchiavelli | Ecology | 1998 | 10.2307/176863 |
| *Khaya senegalensis* | Plantae | Gaoue; Ticktin | Conserv Biol | 2010 | 10.1111/j.1523-1739.2009.01345.x |
| *Magnolia macrophylla var. dealbata* | Plantae | S√°nchez-Vel√°squez; Pineda-L√≥pez | Popul Ecol | 2010 | 10.1007/s10144-009-0161-5 |
| *Manilkara zapota* | Plantae | Cruz-Rodr√≠guez; Lopez-Villavicencio; Valverde | J Trop Ecol | 2009 | 10.1017/S0266467408005713 |
| *Phyllanthus emblica* | Plantae | Ticktin; Ganesan; Paramesha; Setty | J Appl Ecol | 2012 | 10.1111/j.1365-2664.2012.02156.x |
| *Phyllanthus emblica* | Plantae | Ellis; Williams; Lesica; Bell; Bierzychudek; Bowles; Crone; Doak; Ehrl√©n; Ellis-Adam; McEachern; Ganesan; Latham; Luijten; Kaye; Knight; Menges; Morris; den Nijs; Oostermeijer; Quintana-Ascencio; Shelly; Stanley; Thorpe; Ticktin; Valverde; Weekley | Ecology | 2012 | 10.1890/11-1052.1 |
| *Phyllanthus indofischeri* | Plantae | Ticktin; Ganesan; Paramesha; Setty | J Appl Ecol | 2012 | 10.1111/j.1365-2664.2012.02156.x |
| *Pinus lambertiana* | Plantae | van Mantgem; Stephenson | J Ecol | 2005 | 10.1111/j.1365-2745.2005.01007.x |
| *Pinus lambertiana* | Plantae | Maloney; Vogler; Eckert; Jensen; Neale | Forest Ecol Manag | 2011 | 10.1016/j.foreco.2011.05.011 |
| *Pinus nigra* | Plantae | Buckley; Brockerhoff; Langer; Ledgard; North; Rees | J Appl Ecol | 2005 | 10.1111/j.1365-2664.2005.01100.x |
| *Pinus strobus* | Plantae | M√ºnzbergov√°; Hadincov√°; Wild; Kindlmannov√° | PLOS ONE | 2013 | 10.1371/journal.pone.0056953 |
| *Prioria copaifera* | Plantae | Condit | Forest Ecol Manag | 1993 | 10.1016/0378-1127(93)90045-O |
| *Prosopis laevigata* | Plantae | Bernal | NA | 2004 | NA |
| *Prunus africana* | Plantae | Stewart | NA | 2001 | NA |
| *Rhododendron ponticum* | Plantae | Salguero-G√≥mez | NA | 2004 | NA |
| *Rhododendron ponticum* | Plantae | Salguero-G√≥mez | NA | 2004 | NA |
| *Sapium sebiferum* | Plantae | Renne | NA | 2001 | NA |
| *Shorea leprosula* | Plantae | Chen; Visser; Jongejans; van Breugel; Zuidema; Kassim; de Kroon | J Ecol | 2011 | 10.1111/j.1365-2745.2011.01825.x |
| *Styrax obassis* | Plantae | Abe; Nokashizuka; Tanoka | J Veg Sci | 1998 | 10.2307/3237044 |
| *Taxus floridana* | Plantae | Kwit; Horvitz; Platt | Conserv Biol | 2004 | 10.1111/j.1523-1739.2004.00567.x |
| *Tsuga canadensis* | Plantae | Lamar; McGraw | Forest Ecol Manag | 2005 | 10.1016/j.foreco.2005.02.056 |
| *Ziziphus jujuba* | Plantae | Zull; Lawes; Cacho | Environ Modell Softw | 2015 | 10.1016/j.envsoft.2015.10.026 |
| *Oeceoclades maculata* | Plantae | River√≥n-Gir√≥; Ravent√≥s; Damon; Garc√≠a-Gonz√°lez; M√∫jica | Biol Invasions | 2019 | 10.1007/s10530-019-01945-7 |

**Table S2**. Summary of ISO3 country codes used in Figure 1B, GDP, population size (in 2017) and per capita GDP. Source: United Nations.

| **Country name** | **ISO3 code** | **GDP** | **Population** | **GDP per Capita** |
| --- | --- | --- | --- | --- |
| Argentina | ARG | 6.3743E+11 | 43937140 | 14508 |
| Armenia | ARM | 1.1537E+10 | 2944791 | 3918 |
| Australia | AUS | 1.3234E+12 | 24584620 | 53831 |
| Austria | AUT | 4.1684E+11 | 8819901 | 47261 |
| Azerbaijan | AZE | 4.0748E+10 | 9845320 | 4139 |
| Belgium | BEL | 4.9476E+11 | 11419748 | 43325 |
| Belize | BEL | 1862614800 | 375769 | 4957 |
| Belarus | BLR | 5.4456E+10 | 9450231 | 5762 |
| Brazil | BRA | 2.0536E+12 | 207833823 | 9881 |
| Canada | CAN | 1.6471E+12 | 36732095 | 44841 |
| Switzerland | CHE | 6.7897E+11 | 8455804 | 80296 |
| China | CHN | 1.2238E+13 | 1421021791 | 8612 |
| Cyprus | CYP | 2.2054E+10 | 1179678 | 18695 |
| Czech Republic | CZE | 2.1591E+11 | 10641034 | 20291 |
| Germany | DEU | 3.6932E+12 | 82658409 | 44680 |
| Denmark | DNK | 3.2987E+11 | 5732274 | 57545 |
| Spain | ESP | 1.3143E+12 | 46647428 | 28175 |
| Estonia | EST | 2.6612E+10 | 1319390 | 20170 |
| Finland | FIN | 2.523E+11 | 5511371 | 45778 |
| France | FRA | 2.5825E+12 | 64842509 | 39827 |
| United Kingdom | GBR | 2.6379E+12 | 66727461 | 39532 |
| Georgia | GEO | 1.5081E+10 | 4008716 | 3762 |
| Greece | GRC | 2.0309E+11 | 10569450 | 19214 |
| Croatia | HRV | 5.5213E+10 | 4182857 | 13200 |
| Hungary | HUN | 1.3976E+11 | 9729823 | 14364 |
| India | IND | 2.6507E+12 | 1338676785 | 1980 |
| Indonesia | IND | 1.0154E+12 | 264650963 | 3837 |
| Iran | IRA | 4.5401E+11 | 80673883 | 5628 |
| Ireland | IRL | 3.3143E+11 | 4753279 | 69727 |
| Iceland | ISL | 2.4488E+10 | 334393 | 73233 |
| Italy | ITA | 1.9438E+12 | 60673701 | 32038 |
| Japan | JPN | 4.8724E+12 | 127502725 | 38214 |
| Lithuania | LTU | 4.7544E+10 | 2845414 | 16709 |
| Luxembourg | LUX | 6.2316E+10 | 591910 | 105280 |
| Latvia | LVA | 3.0463E+10 | 1951097 | 15613 |
| Moldova | MDA | 8128493432 | 4059684 | 2002 |
| Mexico | MEX | 1.1509E+12 | 124777324 | 9224 |
| North Macedonia | MKD | 1.128E+10 | 2081996 | 5418 |
| Malta | MLT | 1.2518E+10 | 437933 | 28585 |
| Montenegro | MNE | 4844592067 | 627563 | 7720 |
